# Development of an Open-source Low-cost Pressure Myography and Cardiac Flow Simulator, HemoLens, for Mechanical Characterization of Native and Engineered Blood Vessels

**DOI:** 10.1101/2025.04.29.651300

**Authors:** Antonio J. PereiraTavares, Liam Aranda-Michel, Scott Hahn, Brian D. Coffin, Syed Faaz Ashraf, Jason M. Szafron, Adam C. Straub, Daniel J. Shiwarski

## Abstract

Pressure myography, the standard for assessing vascular mechanics and vasoreactivity, is costly ($40,000+), has low throughput, and is limited to static fluid flow. Here, we developed HemoLens, an open-source 3D-printed pressure myography system for ∼$700. HemoLens features compact micromanipulators, incremental in-line pressure control, physiological temperature regulation, and modular pulse pressure control between normotensive and hypertensive levels. HemoLen’s efficacy was demonstrated by delineation of physiological reactivity and pathological mechanical phenotypes using native mouse arteries and bioprinted acellular scaffolds. Wildtype vessels show greater distention (124.3 vs. 43.07 µm) and increased dynamic compliance compared to diseased vessels. Small diameter (450 µm) collagen-based artery-like scaffolds are FRESH bioprinted to mimic hypertensive vascular stiffening. Engineered hypertensive vessels demonstrate increased burst pressure (464 mmHg) and reduced dynamic compliance reminiscent of diseased arteries. Together, HemoLens lowers the barrier to entry in pressure myography research by serving as a comprehensive low-cost system for native and engineered vessel characterization.

**Graphical Abstract:** 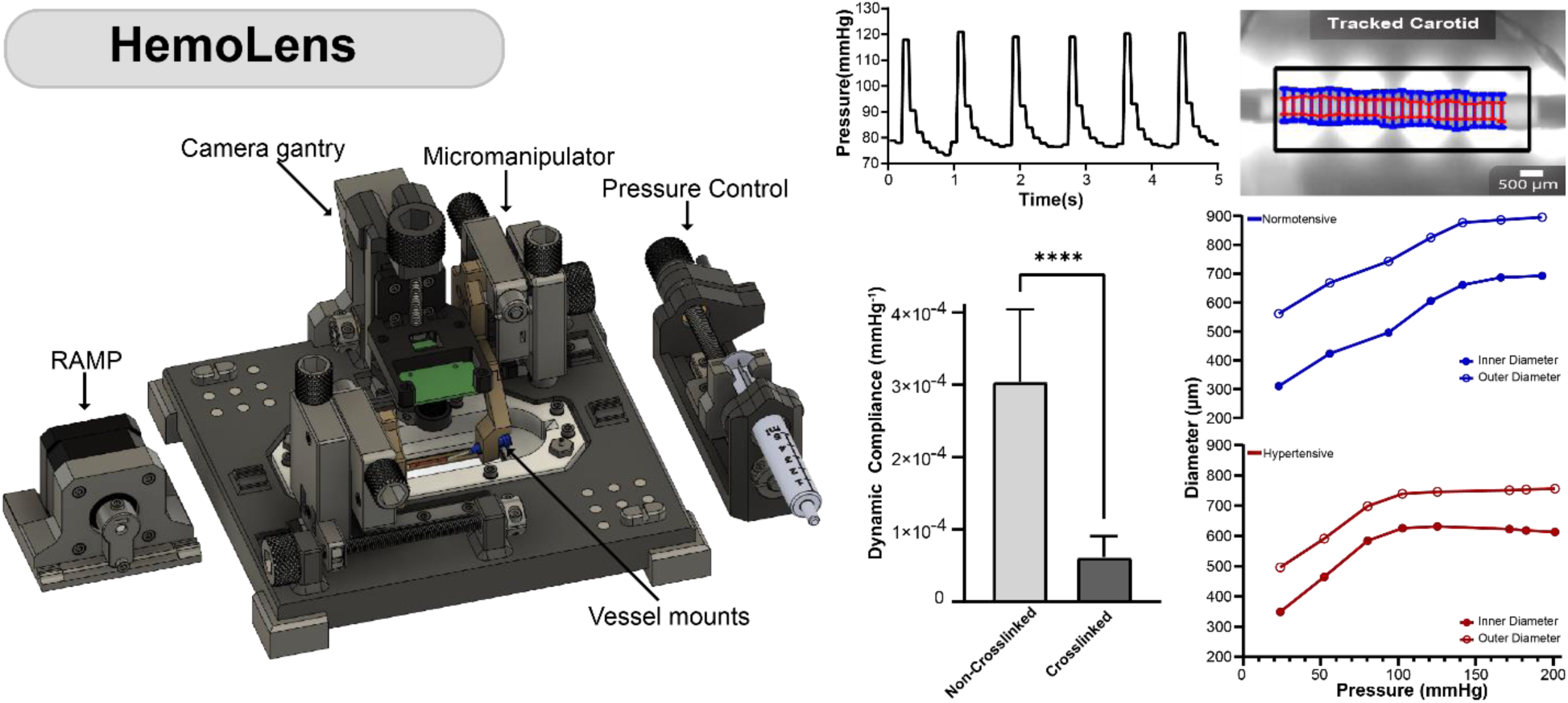

## INTRODUCTION

Arterial vasoreactivity—the ability of small-diameter blood vessels to regulate luminal diameter via constriction and dilation in response to pharmacologic and endogenous stimuli—plays a crucial role in the systemic control of blood pressure^1,2^. These vasoactive blood vessels are unique in that their cellular and extracellular matrix (ECM) components are structured to withstand the unique hemodynamic forces applied by blood flow and pressure changes during the cardiac cycle^3,4^. Given that arterial stiffness is a defining characteristic of hypertension, the resulting increased blood pressure subsequently leads to downstream effects such as endothelial dysfunction and vascular smooth muscle cell de-differentiaion^5,6^. This drives pathological ECM remodeling that promotes systemic vessel stiffening from overproduction of fibrillar collagen, wall thickening, and reduced luminal diameters that limit the vessels’ ability to expand and contract^4^.

Animal models, particularly in mice, have been instrumental in advancing our understanding of hypertension and related cardiovascular conditions. However, these models often fail to accurately replicate human clinical outcomes^7–9^ . As a result, there is growing interest in tissue engineering (TE) approaches, which employ human-specific cells and biomaterials to create biologically relevant models that better mimic human pathophysiology^10,11^. One such approach, 3D bioprinting, enables the precise spatial deposition of ECM-based biomaterials to fabricate a range of tissue-engineered blood vessels (TEBVs)^12^, including acellular vascular conduits^13^, cellularized scaffolds^14^, and perfusable microvascularized functional tissues^15,16^ for both disease modeling and regenerative medicine applications. However, as the physiological relevance of TEBVs continues to improve, there is a growing need for affordable implantation of standardized, reproducible methods to rigorously benchmark their biomechanical properties against native tissues. Moreover, since mechanical dysfunction, such as increased arterial stiffness, is a hallmark of vascular diseases like hypertension, accurate quantification of the mechanical properties of both native and engineered vessels is essential.

Clinical *in-vivo* techniques such as pulse wave velocity and Coronary Flow Velocity Reserve provide information on the mechanical behavior of blood vessels, but do not yield direct insight into the underlying biomechanics of the vessel wall^17,18^. *Ex-vivo* characterization, on the other hand, allows for tightly controlled environments to discriminate between cellular and extracellular remodeling processes^19^. Many traditional mechanical testing methods have been applied to both native and engineered blood vessels to test various mechanical characteristics^20–22^. Yet pressure and wire myography remain the most ubiquitous methods for assessing vascular mechanics and graft compatibility for clinical use by measuring maximum burst pressure and vascular tone^22,23^. Despite their ubiquity, however, commercial myograph systems can cost upwards of $40,000, are limited in throughput, and lack capabilities for customization. Recent open-source myography systems, such as “Vasotracker”, provide excellent software packages for monitoring vessel mechanics and a more affordable hardware platform, but still require custom CNC manufacturing and standalone microscopic imaging systems that drive costs above $6000^24^. Moreover, no current system allows for experimentation in dynamic pressure environments that mimic physiologic cyclic pressure changes. Thus, there is an emerging need for a low-cost biomechanical myography platform specifically designed for testing small diameter (< 1 mm) arteries and resistance vessels that uses affordable manufacturing processes, integrated and accessible imaging systems, and is capable of evaluating both conventional burst pressure and functional vasoreactivity under physiological conditions.

To fulfill this need, we determined several critical components required for a comprehensive system for hemodynamic and material characterization: 1) systemic pressure measurement^25^, 2) optical vessel diameter tracking^26^, 3) a variable intensity light source^24^, 4) accurate micropositioning for easy vessel attachment and tensioning^23^, 5) intraluminal perfusion and media bath circulation at physiologically relevant temperatures^26^, and 6) discrete pressure control^25,27^. Incorporation of these components into one system is challenging and often cost-prohibitive. However, with the hardware and software innovations enabled by the open-source 3D printing and maker community, new devices can now be created with significantly lower costs while maintaining high performance^28^. To this end, we designed a modular open-source pressure myography system built from primarily 3D-printed and low-cost linear motion components called HemoLens. The development and validation of HemoLens was conducted through a multi-step process. First, all mechanical aspects of the system were meticulously characterized. Next, we provide examples of HemoLens’ performance by identifying distinct mechanical differences between normotensive and hypertensive vessels. Specifically, we evaluated burst pressure, dynamic compliance, and vasoreactivity of mouse carotid arteries, recording altered compliance of hypertensive vessels compared to our normotensive control. Lastly, we used HemoLens to examine the mechanical differences between bioengineered Freeform Reversible Embedding of Suspended Hydrogel (FRESH) 3D printed acellular vascular scaffolds as a proof-of-concept towards biofabrication of hypertensive disease models. Taken together, HemoLens is a modular, open-source hardware platform designed for vascular mechanical assessment of both native and engineered vasculature.

## RESULTS

### Vascular Hemodynamic Forces and HemoLens Design Rationale

When evaluating mechanical properties of native or engineered blood vessels the cellular and extracellular matrix structure, composition, and alignment drives function. In small diameter blood vessels such as mesenteric and carotid arteries, a tri-layered vascular wall is specifically organized to provide mechanical strength and function. The outer Adventitial layer is collagen rich providing mechanical reinforcement to prevent vessel rupture and resist changes in intraluminal pressure (Figure 1A,B)^29,30^. The middle Tunica Media layer contains vascular smooth muscle cells (VSMC) and elastic laminins, and other scaffolding proteins that both regulate vascular tone through VSMC contraction/dilation and provide compliance to dampen the mechanical forces produced by cardiac cycle-driven hemodynamic pressure (Figure 1A The innermost layer, Tunica Intima, contains the vascular endothelial cells (EC), which form a selectively permeable barrier along the vascular lumen to sense and respond to changes in blood flow shear stress (Figure 1A,B)^31^. The innermost layer, Tunica Intima, contains the vascular endothelial cells (EC), which form a selectively permeable barrier along the vascular lumen to sense and respond to changes in blood flow shear stress (Figure 1A,B)^3^.

**Figure 1.**
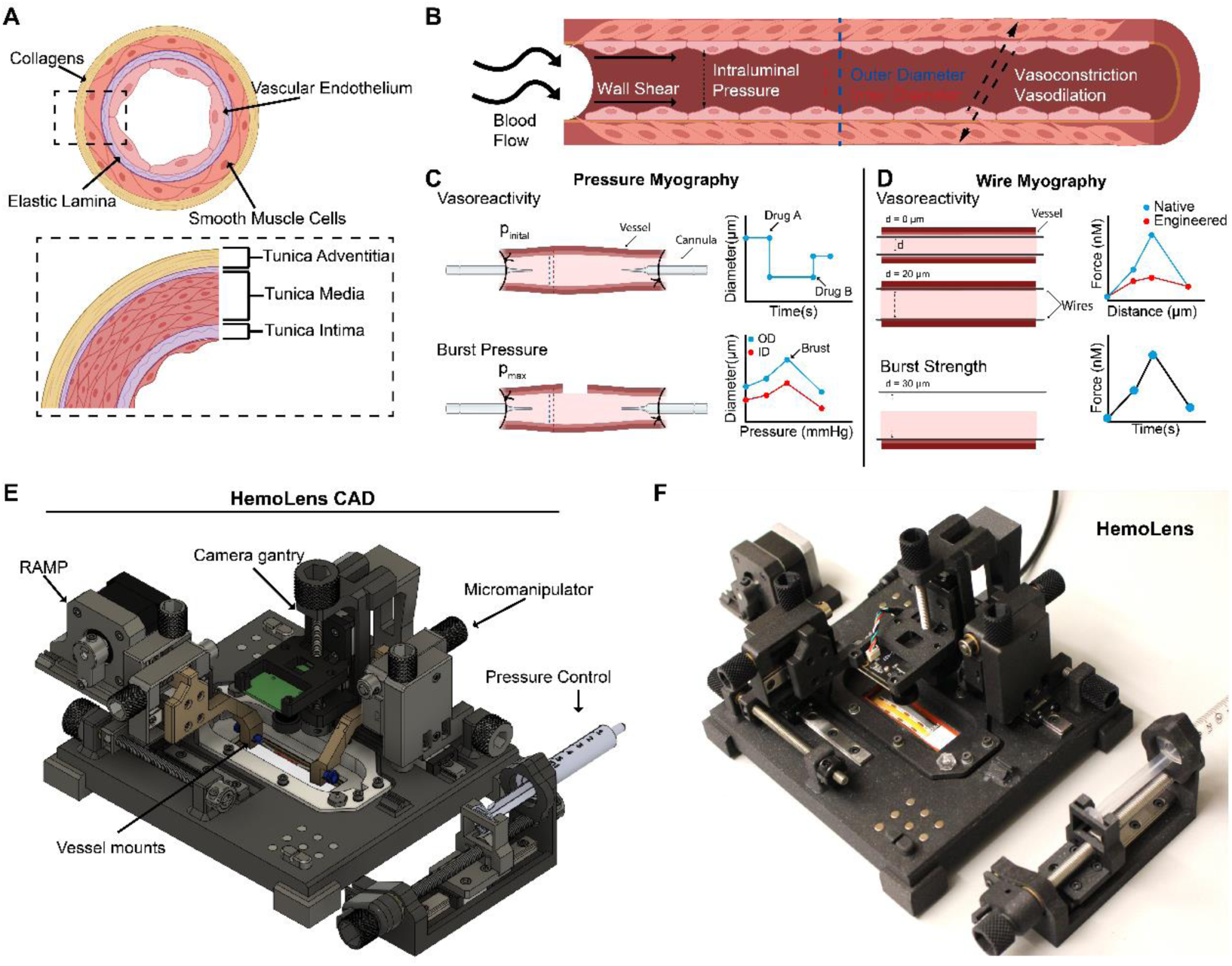
Vascular Hemodynamics and HemoLens Overview **A)** Graphical representation of the three major layers of the blood vessel and their extracellular components. **B)** Graphical representation of the major hemodynamic forces experienced by resistance arteries. **C)** Schematic of Pressure myography showing maximum pressure required to burst a vessel, and representative cartoon graphs of burst pressure testing (i) and myogenic tone (ii). **D)** Wire myography schematic showing the rupture of the blood vessel and representative graphs of rupture testing (i) and myogenic tone (ii) experiments conducted

To isolate the functional consequences of altered vascular biomechanical properties related to developmental and disease progression, researchers perform *ex vivo* characterization with either pressure and/or wire myography. In pressure myography, excised vessels are canulated one at a time, and intraluminal pressure is set to either replicate physiological pressure (120/80 mmHg) and measure vascular tone or increased until the vessel wall ruptures to evaluate structural burst pressure (Figure 1C). Alternatively, in wire myography, an excised vessel is cut into small sections, and two small wires are inserted into the lumen. The wires are then slowly pulled apart until reaching a tensile force that simulates physiological pressure to assess vasoreactivity, or is advanced incrementally until the wires tear through the vessel wall to determine the rupture force (Figure 1D). While wire myography can be higher throughput than pressure myography, pressure myography more accurately recapitulates the fluid mechanics of the vascular system. Additionally, since pressure myography is considered the ISO standard for clinical applications of vascular biomechanics^32^, we choose to design and build HemoLens as a low-cost alternative to commercial pressure myography systems that would fully recapitulate vascular intraluminal pressure and shear stress under physiological environmental conditions (Figure 1 E-F). HemoLens is a comprehensive hemodynamic and material characterization platform that facilitates 1) real-time systemic pressure monitoring, 2) optical vessel diameter tracking, 3) a variable intensity light source, 4) accurate XYZ micropositioning for easy vessel attachment, alignment, and tensioning, 5) intraluminal perfusion and media bath circulation at physiologically relevant temperatures, and 6) discrete static and dynamic pressure control. The overall design incorporated magnetic attachments to enable modular component addition, providing experimental flexibility and customization between native and engineered vessel mechanical testing. As with most open-source hardware design, our goal was also to reduce the cost of the platform without sacrificing performance and reproducibility. In total, HemoLens costs $730 and requires no additional components other than a personal computer with USB input. At this price point, multiple systems can be built and run in parallel to easily scale pressure myography towards a higher throughput methodology.

### Physiological temperature control and perfusion

Temperature is an essential variable for physiological blood vessel function, including vasoreactivity, and preconditioning prior to mechanical testing^25,33,34^. *Ex-vivo* vascular research is typically conducted at physiological temperature (37 °C) via superfusion from an externally heated solution continuously recirculated through the bath chamber by a closed loop pump system^35^. HemoLens’s vessel bath chamber is constructed from a heat-resistant 3D printed insert with a gasketed glass slide compressed in between the bath chamber and the HemoLens base plate to achieve water tightness and visible light illumination (Figure S1). Additionally, HemoLens was designed to allow for independent control of the bath chamber and vessel perfusion while maintaining physiological temperature for both systems. To achieve bath fluid recirculation, two perfusion ports were integrated into the 3D design of the chamber bottom plate to easily connect tubing via Luer Lock coupling with a superfusion process (Figure S1). To avoid direct heating of the bath chamber, which would result in plastic deformation or warping over time, we utilized a low-cost water bath to heat a media reservoir for circulation set at a flow rate of 19 mL/min (Figure S1). Similarly, perfusion within the vessel is achieved using a low volume peristaltic pump with a flow rate of 1 mL/min, and temperature is maintained by submersion of the pump into the heated water bath (Figure 2A).

**Figure 2.**
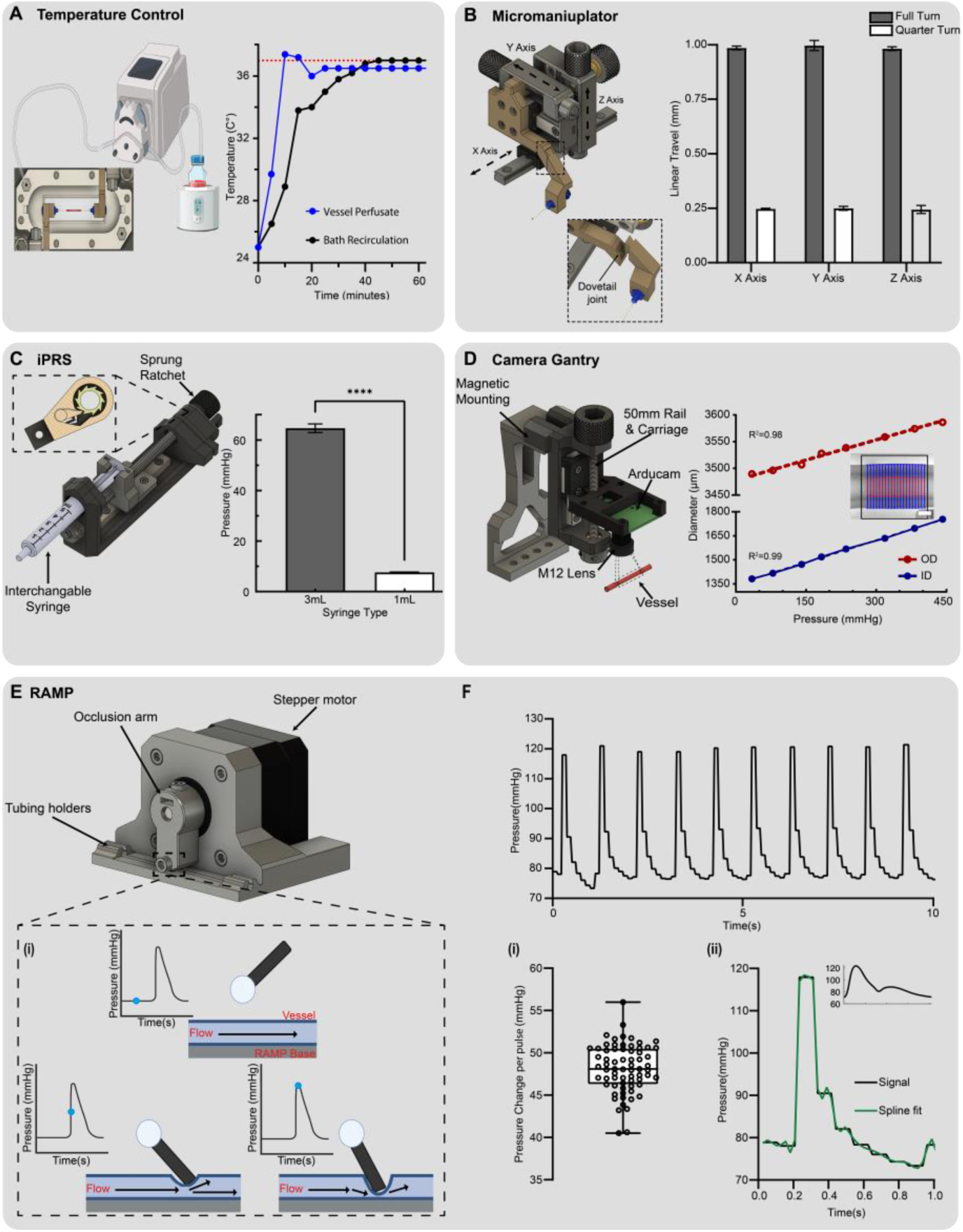
Validation of HemoLens’ components **A)** Temperature reading for both superfusion lines, bath (black), and vessel (grey) at water bath set point of 47C. Both lines reach physiological in 40 minutes and maintain this temperature with a stability of ± 0.32 °C over time. **B)** Micromanipulator highlighting movement axes. (i) Calculated linear movement for each axis. Full Turn: X: 0.985 mm ± 0.009 mm, Y: 0.996 mm ± 0.023 mm, Z: 0.982 ± (0.009). Quarter Turn: X: 0.247 ± 0.003 mm, Y: 0.250 ± 0.009 mm, Z: 0.244 mm ± 0.019 mm. n=10 for all axes. Data are represented as mean ± SEM **C).** incremental pressure ratchet system (iPRS). Inset showing internal stopping mechanisms. (i) measured increase per syringe tested, 3mL: 64.64 mmHg ± 2.94 mmHg, 1mL: 7.62 mmHg ± 0.194 mmHg. n=3 for each syringe types. Data are represented as mean ± SD **D)** Camera gantry system with call outs to important features. (i) Image showing rubber test tube with Vasotracker lines overlayed. (ii) Linear elastic response of the rubber test vessels with incremental pressure increases. R^2^ values of 0.98 and 0.99 for outer and inner diameters, respectively. **E)** RAMP system with call outs to important features. (i) Graphical representation of RAMP’s mechanism of action for cyclic pressure increases. **F)** Representative waveform of RAMP’s cyclic pressure generation set to physiological 80/120 mmHg at 65 beats per minute. (i) Average pressure increases per pulse from the base line, 48.16 mmHg ± 2.85 mmHg, n= 68. (ii) A single isolated square waveform with a spline fit superimposed demonstrating a similar curve to an arterial pulse waveform (inset). Arterial waveform adapted from *Charlton et al 2019*

Defining device equilibration time is critical when designing a vascular characterization platform since excised blood vessels only remain viable and maintain their vasoreactivity for a few hours; even when kept in physiological buffers at 37 °C^25^. To determine the equilibration time to achieve physiological temperature for each perfusion line and longitudinal temperature stability we measured both the circulating bath and vessel temperature for 1 hour. With a water bath temperature set point of 47 °C, HemoLens reached 37 °C within 40 minutes. A stable temperature of 36.63 ± 0.32 °C was maintained over an extended period (Figure 2Ai). With two individually controlled perfusion lines and rapid establishment of physiological temperature, HemoLens offers users quick experimental startup time and multiple avenues of superfusion for the maintenance of physiological temperature.

### High-precision 3D-printed micromanipulators

In all myography systems, precise control of 3-axis positioning is essential to mount vessels onto the testing device to achieve the proper tension without damaging the vessel. Additionally, as the trend toward generating smaller diameter engineered vascular grafts continues, careful and precise manipulation of these vessels will only grow in importance^36,37^. Commercial pressure myography systems employ optical grade manual micromanipulators, but implementation of these components into an open-source device drastically increases the overall cost as they often can be the single most expensive component costing > $1000. Therefore, to dramatically lower the cost of HemoLens we designed and built a compact 3-axis (X, Y, Z) manual micromanipulator for ≤ $50 per unit by leveraging widely available miniature linear rails and carriages from the open-source 3D printing community (Figure 2B). In addition to 3-axis positioning, our micromanipulator design includes a customizable dovetailed arm for standard Luer Lock Birmingham needle gauges to which the vessel is sutured (Figure 2B).

Commercial 3-axis micromanipulators report precision as linear travel per revolution. We employed the same metric to analyze the precision and repeatability for each of the 3 axes of HemoLens’ custom micromanipulators. A complete turn of each axis resulted in 0.99 ± 0.016 mm of average linear travel per axis, and a quarter turn, calculated similarly, resulted in 0.25 ± 0.011 mm of linear travel (Figure 2Bi). These experimental values closely match the expected 1 mm pitch of the M6 bolts used as axis leadscrews suggesting that there is minimal mechanical backlash and high repeatability. With these results, we highlight our manipulator as a low-cost, high-performance, and reliable alternative to commercial 3-axis micromanipulator stages well suited for vessel cannulation and myography applications.

### In line pressure monitoring and discrete pressure control with a designed “ratchet”

Control and monitoring of intraluminal pressure is the critical feature of pressure myography that separates it from other vasoactive and mechanical characterization techniques^23,32^. Many complex systems are used in commercial myography systems to both monitor and automate pressure control; however, these engineered controls are expensive and are normally restricted to sub-burst pressure levels during experimentation. Moreover, since static pressure is often set at the start of an experiment and held constant throughout, we wanted to provide users with a simple, modular, and non-motorized syringe-based pressure control system. To achieve real-time pressure monitoring we implemented a previously developed open-source pressure monitoring system from Vasotracker^24^. Two in-line pressure sensors were attached to the vessel perfusion line before and after the vessel cannulation arms. The range and sensitivity of the flow-through pressure sensors can be tuned to measure small changes in physiologic vasoreactivity or expanded by adjusting the Wheatstone bridge and software configuration to characterize mechanical burst pressure. Following calibration of the pressure sensors, repeatable pressure monitoring within ± 3.43 mmHg was achieved within the working range of the pressure sensors (≤ 500 mmHg ) (Figure S1).

Next, the HemoLens incremental pressure ratchet system (iPRS) was designed to function as a low-cost manually advancing syringe pump (Figure 2C). In a closed-loop manual pressure control system, as pressure rises, back pressure accumulates at the pressure-generating site. This pressure increase can push the syringe plunger backward, ultimately reducing the overall systemic pressure. To avoid this backpressure driven pressure loss during experimentation, we designed an internal anti-slip rotatory “ratchet” mechanism to prevent anti-clockwise movement when vessels are pressurized (Figure 2C insert). A manual clutch disengagement system was also incorporated at the top of the ratchet mechanism to allow easy syringe retraction to the starting position or to quickly depressurize the vessel. HemoLens’ iPRS ratcheting pressure control module is fully customizable, enabling users to select various syringe diameters for incremental pressure tuning.

To characterize the incremental pressure repeatability across a range of syringe diameters we measured discrete pressure intervals for 3 mL (8.66 mm ID) and 1 mL (4.78 mm ID) BD syringe. Rubber tubing was mounted and sutured onto HemoLens as a minimally compliant synthetic test vessel, and pressure was increased in a stepwise manner by turning the lead screw on the iPRS. Each counterclockwise turn could be audibly perceived as a “click” to provide user feedback in response to incremental advances. Each click increased pressure by 64.37 ± 2.95 mmHg for 3 mL syringes and 7.62 ± 0.19 mmHg for 1 mL syringes. (Figure 2C-i). While this difference in pressure exceeds the theoretical ∼4.6X change for an adiabatic process between these syringes, the variations in plastic syringe plunger style and overall syringe rigidity likely influence the experimentally observed pressure change. With this iPRS system, we demonstrate both high precision and broad range control over pressure increases in a low-cost, compact design that does not require motorization.

### Modular low-cost optical vessel tracking

Most myography devices employed for investigating vascular reactivity and mechanical properties utilize optical imaging of the inner diameter (ID) and outer diameter (OD) of the mounted vessel to compute changes in diameter, wall thickness, and compliance. To perform this function, commercial and open-source systems conventionally rely on external microscopy systems with in-line lighting solutions, which increases overall system cost and limits throughput based upon optical system avalibility^19^. Our objective was to eliminate optical imaging as a hardware constraint by utilizing new advances and price reductions in CMOS sensors and LED lighting to achieve integrated, low-cost, high-throughput imaging. We first designed a custom imaging gantry for an ArduCam USB camera with an M12 ArduCam Lens (Figure 2D). The camera mount was then attached to a 50 mm linear rail for precise control of camera focus. A magnetic camera holder was designed to easily attach and remove the camera gantry from HemoLens’s base. (Figure 2D). This removable design improved the vessel mounting procedure by eliminating any overhead obstruction during alignment and suturing. A variable intensity lighting system was incorporated into the HemoLens base for oblique or in-line lighting (Figure S1). Finally, to ensure HemoLens’ reliability and performance as an optical pressure myography system, compliant rubber tubing (ID of ∼1.4 mm, OD of ∼3.5 mm) was mounted and optically tracked using the open-source Vasotracker software (Figure 2D_i_). A stepwise pressure increase was applied via the manual iPRS pump while monitoring vessel ID, OD, and inlet and outlet pressure (Figure S1). The test synthetic vessel’s diameter increased linearly (R^2^ values of 0.98 ID and 0.98 OD) as a function of pressure up to 450 mmHg (Figure 2D_ii_). These results demonstrate HemoLens’ capability to conduct conventional real-time pressure myography with a low-cost and modular CMOS sensor.

### Cyclic physiological pulsatile pressure generated with RAMP

Despite their ubiquity and relevance, state-of-the-art ex-vivo vascular research platforms often rely solely on static pressure to measure vessel reactivity and burst pressure^20^. This neglects the dynamic native vessel environment, characterized by cyclic pulsatile flow that generates pressure waves between 80 mmHg (diastolic) and 120 mmHg (systolic) in normotensive adult humans. In hypertensive pathological conditions, this flow can exceed 145 mmHg systolic pressure^1,38^. A low-cost aquarium pump was used to establish closed-loop luminal flow through our cannulated test vessel. Systemic baseline pressure was achieved by attaching a custom 3D-printed occlusion collar to the outlet tubing and manually restricting flow until physiological 80 mmHg was achieved. To incorporate pulsatile pressure into HemoLens, the Regular Adjustment of Modulated Pressure (RAMP) system was designed (Figure 2E). RAMP consists of a NEMA 17 stepper motor attached to a 3D printed bracket and occlusion arm that partially occludes the outlet tubing line via a swinging motion to create a brief pulsed increase in systemic pressure that matches the physiological arterial pressure waveform (Figure 2E_i_). RAMP’s occlusion arm can be altered by adjusting the distance between the occlusion arm and outlet tubing to tune the maximum peak pressure (Figure 2E_i_).

To confirm the repeatability and physiological accuracy of the RAMP system, pulsatile pressure readings were measured using a similar synthetic test vessel as above. When the outlet collar was cinched to achieve a baseline of 80 mmHg, RAMP was programmed to create 65 beats per minute or 1.08Hz cyclic pressure increases from ∼80 to ∼120 mmHg. Our results show an average of 48.16 ± 2.85 mmHg pressure increase with each pulse (Figure F_i-ii_). The measured frequency of the RAMP pulse wave sequence was empirically determined to be ∼62 beats per minute (bpm) or ∼1.03 Hz, which is within the frequency range to mimic a normal human heartbeat^39,40^. The slight deviation between our programmed beat rate and measured frequency is likely due to a brief delay in occlusion arm rotation during tubing compression. For RAMP, beat rate is easily tunable to achieve a desired frequency by altering HemoLens’ C++ code, as is the maximum pulsatile pressure by adjusting RAMP’s occlusion arm length or adding a spacer beneath the perfusion tube to increase tube compression. Additionally, as both the rate and degree of occlusion can be tuned, researchers can use RAMP to mimic physiological and pathological pressure conditions, effectively incorporating human heartbeat pulsatile pressure into their vascular research.

### Wildtype and Diseased Vessel Characterization with HemoLens

The pathological hallmark of hypertensive vessel stiffening is the overproduction of collagen and degradation of the elastin of the vessel wall following damage to the endothelium^41^. The stiffer a vessel becomes, the more impaired its ability to dilate and contract to accommodate fluctuations in blood flow and pressure^42^. To demonstrate HemoLens’ capability to elucidate static and dynamic responses to pressure changes in diseased vessels, bilateral carotid arteries were excised from both wildtype (WT) and Sickle Cell Anemic (SCA) mice, which are known to exhibit endothelial damage-associated vascular remodeling and vessel stiffening^43,44^ (Figure 3A). In this pilot study, there were very little differences between the inner and outer diameters of WT and SCA carotid arteries when measured by stereomicroscopy (ID: 314.98 ± 17.75 µm vs. 328.29 ± 38.55 µm and OD: 439.83 ± 16.40 µm vs. 530.58 ± 22.9 µm, respectively) (Figure S2). Each excised mouse artery was mounted onto HemoLens for evaluation (Figure 3B). To ensure the mounted vessels were vasoreactive post excision, excised WT mouse carotid arteries were exposed to increasing concentrations of Phenylephrine (PE) and Acetylcholine (ACh) as previously described^45,46^. A noticeable increase in constriction was observed at 10^-6^ M Pe, reaching a maximum constriction at 10^-5^ M PE (Figure 3C). When exposed to ACh, vessel relaxation increased linearly with increasing concentrations of ACh (Figure 3D). The highest range of the working concentration of ACh was deemed the point of maximum relaxation. However, true maximum relaxation experiments are beyond the scope of this work. Since maintenance of vasoreactivity requires a stable physiological experimental environment (temperature, salt concentrations, and pressure) to ensure experimental repeatability^47^, these results highlight HemoLens’ ability to control critical environmental conditions during myography experimentation.

**Figure 3.**
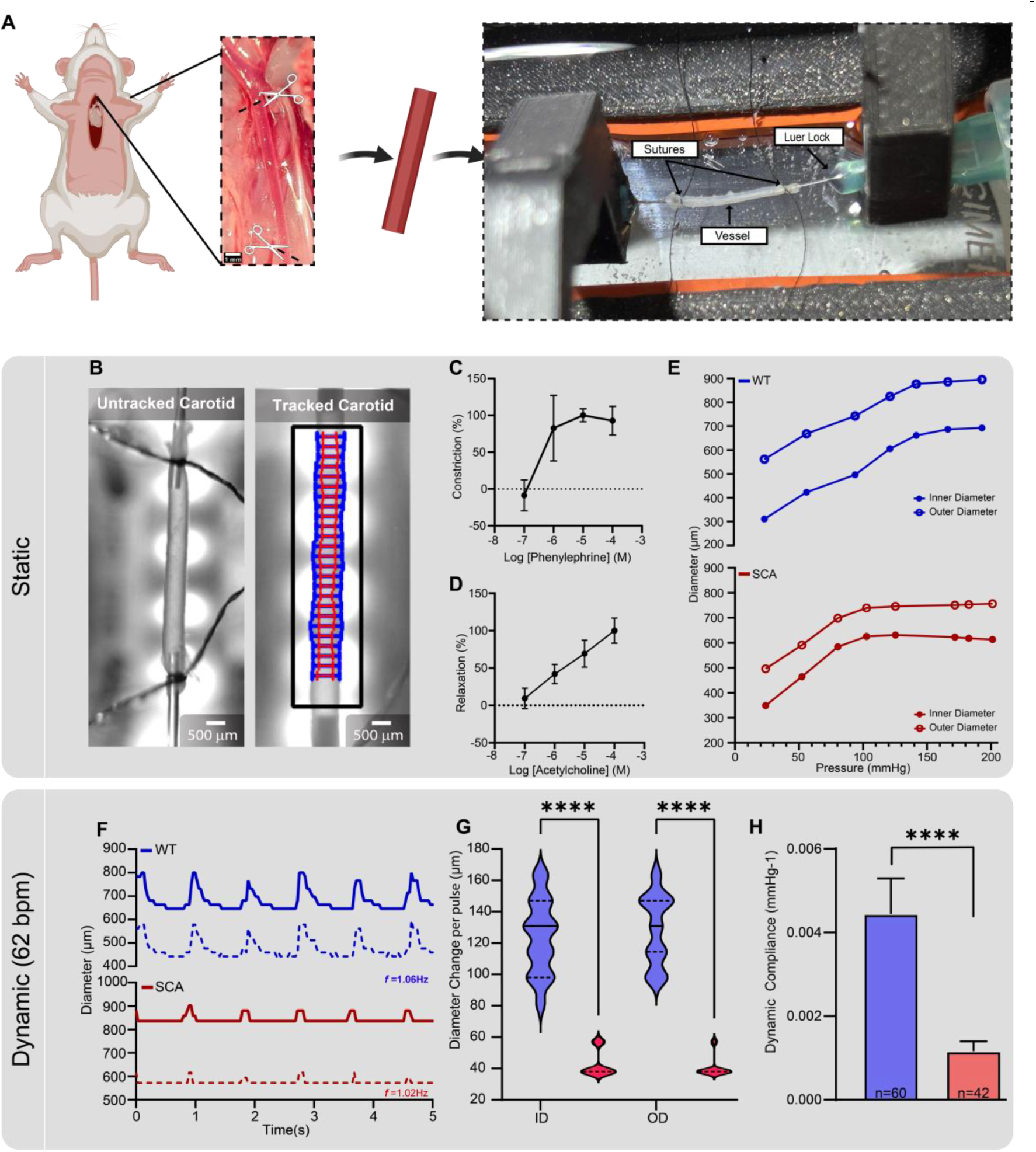
Native artery characterization with HemoLens **A)** Schematic showing carotid artery excision from mouse and mounted on HemoLens. **B)** Images of mounted carotid arteries with and without Vasotracker trace lines overlayed. **C)** Normalized constriction percentage of WT artery exposed to cumulative concentrations of Phenylephrine with maximum constriction occurring at 10^-5^ M. Constriction was measured using outer diameter **D)** Normalized vessel relaxation percentage of WT artery exposed to cumulative concentrations of Acetylcholine. No maximum relaxation was recorded. **E)** Pressure myography results demonstrating a lower elastic response to internal pressure increases of SCA vessels (red) reach peak dilation at 150 mmHg, while WT vessels (blue) show a peak dilation at 200 mmHg. **F)** Representative wave forms of WT (blue) and SCA (red) vessels under physiological RAMP generated pressure waves at 62 beats per minute. **G)** Average diameter change per pulse in each vessel. WT vessels how higher changes per pulse for both inner and outer diameters: 124.3 ± 23.59 mmHg (n=63) vs. 43.07 ± 8.49 mmHg (n=45), 129 ± 22.19 mmHg (n=63) vs. 40.62 ± 6.61 (n=48) mmHg, respectively. **H)** Calculated dynamic compliance of each vessel under physiological RAMP generate pressure waves. WT vessels (0.0045 ± 0.0008 mmHg^-1^, n= 61) are significantly more dynamically compliant thanSCA vessels (0.0012 ± 0.0002 mmHg^-1^, n= 42). Data are represented as mean ± SD

Stepped intraluminal pressure increase is considered a gold standard for evaluating vascular mechanics such as strength, burst pressure, and compliance of excised and engineered vessels^32^. Therefore, we conducted mechanical testing on our excised mouse vessels using HemoLens as previously described^23^. The intraluminal pressure of the mounted vessels was increased stepwise with the iPRS pump and held static for 20 seconds at each pressure increment. Both WT and SCA vessels displayed an increase in inner and outer diameter as a function of increasing intraluminal pressure (Figure 3E). Interestingly, the SCA vessel’s diameter-pressure response curve plateaued early at 125 mmHg of internal pressure, whereas normotensive vessels showed higher distensibility and plateaued at 200 mmHg of internal pressure (Figure 3E). Vessel diameters remained unchanged beyond 200 mmHg for both conditions (Figure S2).

### Wildtype vessels have higher dynamic compliance than diseased vessels

Blood flow within the cardiovascular system is dynamic. As blood pressure and heart rate vary cyclically, a vessels diameter distends and contracts in response to these hemodynamic demands^48,49^. This is a phenomenon known as dynamic compliance (DC)^50^. RAMP was specifically designed to generate cyclic pressure waves at human cardiac frequencies, facilitating dynamic compliance analysis of a diverse range of vasculature. The dynamic compliance of mouse carotid arteries was evaluated under physiological pressure variations induced by RAMP. WT vessels showed a much higher average distention of 124.3 ± 23.59 µm for outer diameter and 129 ± 22.19 µm for inner diameter (Video S1, S2). In contrast, SCA vessels only dilated to an average of 43.07 ± 8.50 µm OD and 40.62 ± 6.61 µm ID (Figure 3F-G). The pulse frequency in these native vessels was found to be 1.06Hz and 1.02Hz for WT and SCA, respectively. A comparison of the calculated dynamic compliance is shown in Figure 3H. The calculated dynamic compliance for carotid arteries during pulsatile flow recorded over one minute is significantly higher in WT vessels (4.445^-3^) compared to SCA vessels (1.160^-3^). This outcome, in conjunction with the incremental pressure results, underscores HemoLens’s capability to distinguish various disease relevant mechanical phenotypes. Furthermore, it enables the observation of the inherent pathological vascular stiffening under pulsatile pressure associated with SCA.

### FRESH bioprinting of collagen vascular scaffolds

3D bioprinting is a biofabrication method that has gained immense popularity in recent years^51,52^. With the ability to create patient specific geometry and control scaffold microstructure^53^ (Hudson paper), it is no surprise that many researchers have begun to employ bioprinting as a fabrication method^54–56^. To demonstrate HemoLens’ applicability beyond native vessels and its usefulness in biofabrication workflows, a small-diameter collagen vascular scaffold was designed as a proof-of-concept bioprinted test vessel. To increase experimental throughput and improve printability, we designed a single scaffold model containing five free-floating artery-like vessels supported within a collagen frame. Each individual vessel was designed to have a 450 µm inner diameter and a 1200 µm outer diameter (Figure 4A_i_). The model was sliced into individual layers for machine pathing, visualized to confirm appropriate model settings, and exported as 3D printer Gcode (Figure 4A_ii_).

**Figure 4.**
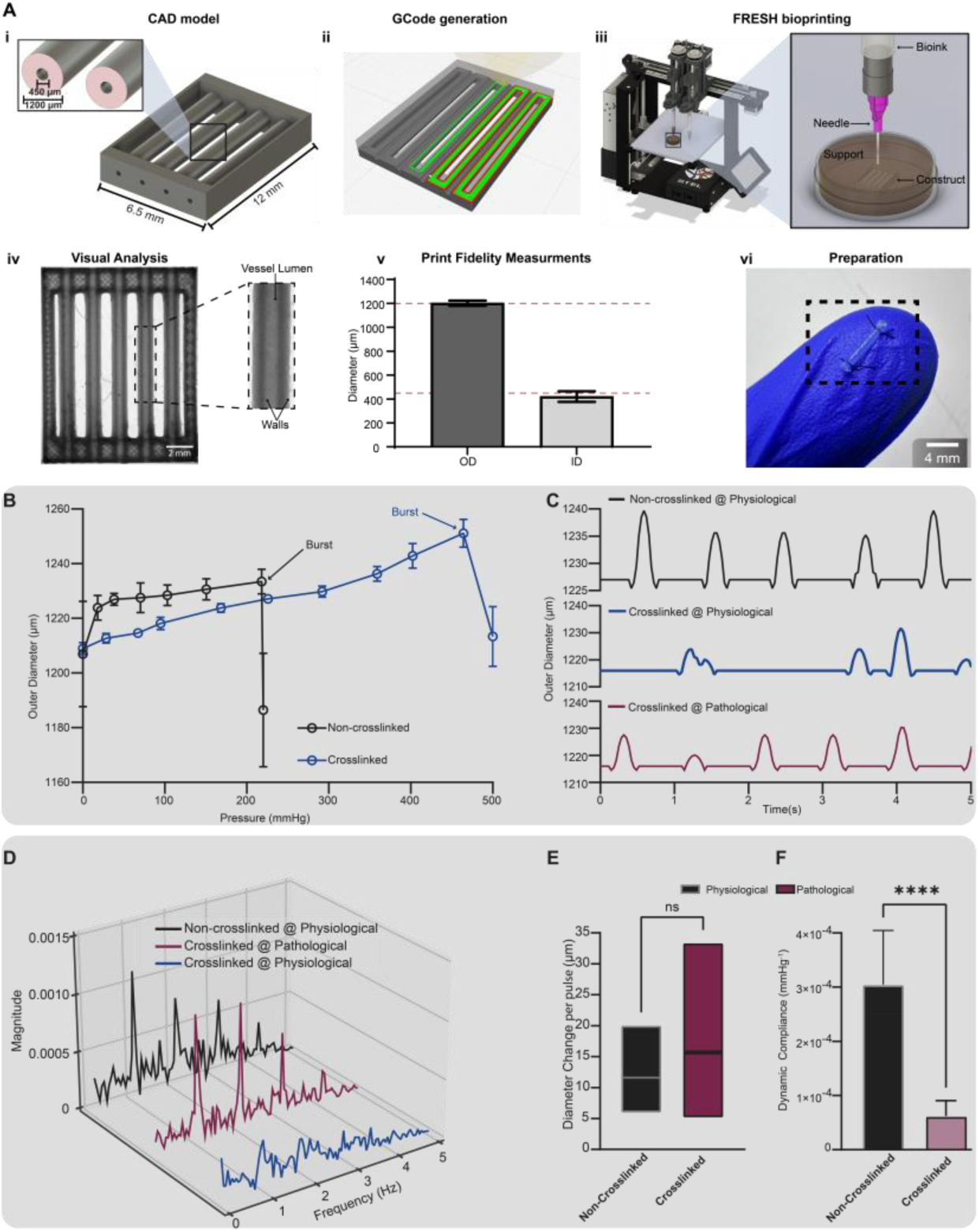
FRESH printed scaffolds evaluated with HemoLens. **A)** Simplified workflow of FRESH biorprinting of test scaffolds. (i) CAD model showing designed 450 µm ID and 1200 µm OD. (ii) Visualized GCODE/Machine pathing using Cura v5.4. (iii) Graphical representation of FRESH bioprinting. (iv) Stereomicroscope image of FRESH printed test scaffold array. SB= 1000 µm. (v) Post processing measurements of the printed scaffold using the measurement tool in LasX v3.9 (Leica). OD = 1172 ± 25.88 µm (n=5), ID = 430 ± 11.88 µm (n=5). All prints used were within a 5% deviation from the original design. (vi) Photo showing excised test vessel during HemoLens mounting step. **B)**. Burst pressure testing of non-crosslinked and 4% PFA crosslinked scaffolds. Crosslinked scaffolds had nearly double the burst pressure of non-crosslinked vessels, 464.34 mmHg and 218.18 mmHg, respectively. **C**) Representative waveforms of each scaffold condition under RAMP generated pressure waves. Crosslinked vessels (blue) do not show a regular diameter change pattern in response to physiological pressure waves generated by RAMP. When under pathological pressure conditions (80/180+ mmHg) crosslinked vessels (red) show a similar pattern to non-crosslinked (black) at physiological (80/120 mmHg) conditions. **D)** Spectra graph of a Fast Fourier Transform (FFT) on the wave forms of C showing that crosslinked at pathological (red) and non-crosslinked at physiological (black) have similar frequencies, while no uniform frequency is seen with crosslinked scaffolds at physiological conditions (blue). **E)** Calculated diameters change per pulse for non-crosslinked under physiological conditions and crosslinked under pathological conditions are shown to be non-significantly different. 11.60 ± 3.57 µm vs. 15.67 ± 7.335 µm. **F)** Calculated dynamic compliance for non-crosslinked under physiological conditions (3^-4^ ± 9.9^-5^ mmHg^-1^, n= 140) is significantly more dynamically compliant and crosslinked scaffolds under pathological conditions (6.23^-5^ ± 2.83^-5^mmHg^-1^, n= 92). Data are represented as mean ± SD.

To create small-diameter artery-like vascular scaffolds, we utilized the FRESH 3D bioprinting process. FRESH enables the accurate fabrication of collagen-based scaffolds by extruding the biomaterial into a gelatin microparticle support bath^57^. The support bath contains a pH buffer to neutralize the acidic collagen biomaterial, resulting in immediate gelation as the layers are printed. Once the printing process is complete, the temperature of the bath is raised to 37°C allowing for non-destructive print release and retrieval^58^.

Each vascular scaffold was FRESH as previously described^59^ from 70mg/mL of collagen-I. The printed scaffolds exhibit a patent luminal morphology characterized by a clear distinction between the transparent lumen region surrounded by a thick opaque (darker in image) outer wall (Figure 4A_iii-iv_). Quantification of the vessel’s inner and outer diameters confirmed that our scaffolds are within a 5% deviation of the original design. The outer diameter was measured to be 1202 ± 21.60 µm, while the inner diameter was 420.6 ± 44.07 µm (Figure 4A_v_). Prior to experimentation, each individual vessel was dissected from the collagen frame and prepared for myography quantification.

### Crosslinking Alters Vessel Material Properties Toward a Stiffened Vascular Phenotype

Due to its excellent biocompatibility, collagen is one of the most widely used biomaterials in tissue engineering^60,61^. However, scaffolds made from collagen alone are notoriously weak and often require post processing modifications, such as chemical crosslinking, to increase overall material strength and suturability^62^. We hypothesized that crosslinking would enhance the printed scaffold’s mechanical strength, but it would potentially compromise the dynamic compliance, resembling a stiffened disease phenotype. To test this, our printed vessels were crosslinked with 4% (v/v) paraformaldehyde (PFA). Crosslinking significantly increased FRESH printed collagen scaffolds’ burst pressure, resulting in a doubling from 218 mmHg to a peak of 464 mmHg (Figure 4B). Scaffolds were then tested at physiologically relevant dynamic pulsatile pressures produced by RAMP. At a physiological rate, control non-crosslinked vessels showed a consistent pulsatile beat frequency of 1.02 Hz (Video S3); however, crosslinked vessels displayed a weak and nonuniform sporadic beat frequency with reduced peak height (Figure 4C, D, Video S4). Interestingly, after switching the crosslinked scaffold to a pathological dynamic pressure regime (peak systolic pressure of 180 + mmHg), we regained a consistent pulsatile beat frequency profile nearly identical to the non-crosslinked scaffolds under physiological pressure (Figure 4C-D, Video S5). The average diameter change per pulse was negligible between non-crosslinked scaffolds at the physiological rate and crosslinked scaffolds at the pathological rate (Figure 4E). Additionally, since a higher pulsatile pressure was required to achieve a similar vascular wall distention, the calculated dynamic compliance of our crosslinked vessel was significantly lower than the non-crosslinked vessels, 6.23^-5^ and 3.04^-4^ , respectively (Figure 4F). Taken together, these data show that with HemoLens we can characterize mechanical phenotypical differences between altered engineered vascular scaffolds and that by implementing FRESH bioprinting, we can mimic the loss of dynamic compliance observed in vascular disease progression.

### Computational Characterization of Wildtype and Disease Vascular Material Properties

Prior work by the Humphrey and Mecham Labs have led to constitutive models for vascular biomechanics describing the non-linear mechanical behavior of arterial vessels^63,64^. In particular, aortic vessels demonstrate a stress vs stretch response curve containing three zones of interest (Figure 5A)^63^. The first is a decreased incremental elastic modulus at low stretch when elastin dominates mechanical behavior, followed by a sharp increase in modulus at high stretch when collagen dominates the behavior. The intersection between these two regions is termed the physiological region^63^. A balance between these mechanical properties is essential to maintain adequate compliance during pulsatile blood flow and resist vascular damage under high pressure.

**Figure 5.**
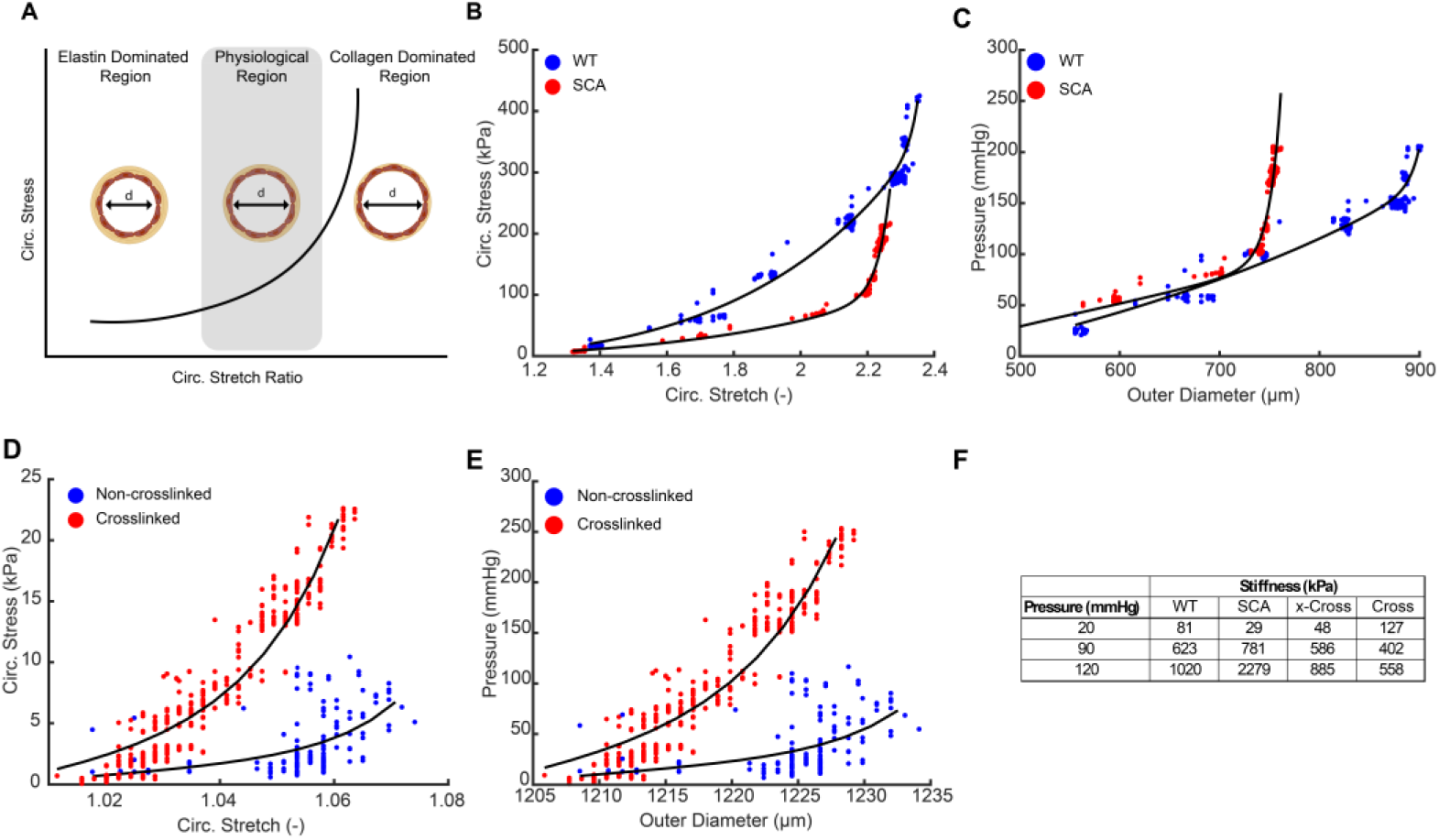
Advance Mechanical Analysis of Vessels and Scaffolds Tested on HemoLens. **A)** Schematic showing non-linear behavior of arterial vasculature. Adapted from *Wagenseil et al 2009*. **B)**. Circumferential stretch and stress curve relationship highlighting the increase compliance of WT vessels within the collagen dominate region denoted by the leftward shift of the SCA vessels. **C)** Pressure diameter relationship of WT and SCA mice displaying characteristics of material stiffening in SCA mouse vessels. **D)**. Circumferential stretch and stress curve relationships between FRESH printed acellular scaffolds. The leftward shift in the stress vs stretch curve is indicative of stiffening at comparable levels of stretch. **E)** Pressure diameter relationship of FRESH printed acellular scaffolds displaying characteristics of increased material stiffness of crosslinked vessels. **F)** Summary table of calculated material stiffnesses at various pressure ranges.

However, during vascular disease, as vessels become less elastic and stiffen due to increased collagen deposition, a shift in the slopes of these regions is observed^63,65^. Therefore, to demonstrate the utility of HemoLens beyond standard diameter tracking, we implemented advanced computational analysis for fitting material parameters to the myography data. We began by extracting material and structural stiffness values using applied techniques for native blood vessel mechanical analysis^66^. Circumferential stress and stretch were calculated for each native sample, showing a much higher collagen dominant region in the SCA mouse vessel compared to the WT mouse vessel, highlighted by the sharp increase in circumferential stress at higher stretch ratios (Figure 5B). Similarly, the calculated material stiffness for three different loading states based on fits to the raw pressure vs diameter data (Figure 5C) and the calculated circumferential stress vs. strain behavior show an increased resistance to diameter expansion in the SCA condition compared to the WT condition. At low pressures (20 mmHg), stiffness is slightly higher in the WT sample, but at a normotensive value of pressure, stiffness is elevated in the SCA sample. At hypertensive pressure (120 mmHg), the SCA sample’s stiffness is substantially elevated compared to the WT sample, as reflected by the sharp increase in slope of the stress vs stretch plot at higher values of stretch. Our advanced computational analysis suggests that the vascular mechanics of our SCA vessels are more collagen dominated than the WT control vessels, which aligns with the known vascular stiffening associated with sickle cell anemia^43^.

We also examined the material stiffness of our printed vessels treated with and without PFA crosslinking agent. As expected, there was a leftward shift in the stress vs stretch curve that was indicative of stiffening at comparable levels of stretch (Figure 5D). However, at comparable pressures, the degree of stretch was much less in the cross-linked samples (Figure 5E), which resulted in lower values of material stiffness for pressures of 90 and 120 mmHg (Figure 5F). Material stiffness at low pressure (20 mmHg) was elevated with crosslinking to the highest value of all samples tested, which appeared consistent with the decreased non-linearity of its response (Figure 5F). Taken together, these advanced biomechanical analyses highlight the broad utility of an open-source HemoLens platform, and serve to guide material property selection towards creating bioprinted acellular scaffolds that can mimic vascular disease progression.

## Discussion

Tissue-engineered vascular scaffolds and Tissue-engineered blood vessels (TEBVs) must meet the mechanical requirements of the native vascular environment before they can be considered for clinical applications^3^. Therefore, mechanical characterization becomes a key metric in developing TEBVs, with intraluminal burst pressure being considered a gold standard^32^. In addition to burst pressure, these scaffolds must remain elastic to maintain their dynamic compliance, expanding and contracting with internal pressure changes to regulate blood flow and maintain natural blood pressure with each pressure cycle. In contrast to the immediate elastic changes captured by the dynamic compliance during a pressure cycle, vasoreactivity involves a more gradual process of smooth muscle contraction and relaxation, altering vessel diameter over extended periods of time, and this phenomenon of vasoreactivity is mainly found within vessels with small luminal diameters (≤ 450 µm)^67,68^. To our knowledge, no comprehensive platform integrates burst pressure myography and cyclic pressure waves into a modular, open-source, inexpensive format. To achieve this, we needed a system that integrates micron-level micromanipulation, an open-source imaging platform, heated luminal perfusion and bath circulation, and a tunable cyclic pulsatile pressure wave. Thus, we developed HemoLens as a bespoke platform for the mechanical assessment and characterization of native vasculature and TEBVs.

Commercial vascular characterization platforms are considered premier equipment in vascular research. However, these systems are expensive and do not test vessels within their native pulsatile environment. Recent open-source myography systems, such as Vasotracker, have significantly reduced the costs of entry into vascular research and provided valuable open-source resources to the community, yet neglect cyclic pressure and are still costly (≥ $6000)^24^. HemoLens addresses this issue by offering pressure myography, cyclic pressure, and physiologically relevant heating in an open-source, inexpensive format. As a myography platform, HemoLens offers micromanipulation down to 250 µm, which rivals commercial systems in performance at 12% of the cost of open-source alternatives and 3% of the cost of commercial systems. The modular imaging system efficiently tracks vessel diameter during experimentation, is magnetically removable, and works across a range of vessel sizes. Additionally, the iPRS manual pressure pump design includes a 3D printed anti-slip clutch that provides consistent and stable discrete pressure increases. However, as a low-cost system, HemoLens balances both performance and cost. For instance, switching from a plastic BD syringe used in this report to a glass Hamilton brand syringe should provide a more linear relationship between syringe size and incremental pressure steps.

Given that most components are fabricated from 3D-printed plastic, it is anticipated that some material wear will occur, particularly in the locking spring of the iPRS ratchet mechanism. However, since HemoLens was designed for modularity and customization, researchers can address these limitations by modifying part design or simply by changing the plastic filament to more durable materials such as polycarbonate or carbon fiber reinforced PETG^69–71^. Additionally, some of HemoLens’ design choices prioritize cost reduction at the expense of potential ease of use. The iPRS ratchet system, for example, requires manual control, limiting experimental automation. Motorization of the manual iPRS pressure control can be achieved by attaching a stepper motor to the leadscrew driving the ratchet system. These types of modifications can be fully integrated into the electronics firmware and added to the Vasotracker open-source software to provide a fully automated experience. Thanks to the maker space community, NEMA 17 stepper motors have become quite inexpensive and typically come with their own Arduino interface cables; thus, an extra stepper motor would have a negligible impact on the overall cost of the HemoLens system. If researchers require higher precision micromanipulation, the resolution of our micromanipulators can be increased by employing higher quality lead screws with a finer thread pitch. To enhance the capabilities of HemoLens, future modifications can be implemented to incorporate load cell force sensors into either of the micromanipulators. This will enable the provision of force readouts, facilitating tensile strength testing, suture retention assessments, and the recording of tensile stress during myography of mounted vessels. Our designed RAMP system provides cyclic pressure increase in the physiological range of 80 - 120 mmHg. The empirically measured 62 bpm was lower than our target of 65 bpm. We believe that further fine tuning of the voltage to the motor or the use of more compliant tubing to reduce the possible drag experienced by the occlusion arm while in contact with the tubing could reach the target bpm.

Mouse models remain a major benchmark in vascular tissue engineering, with native mouse vessels used as the control group for many in-vivo studies^72^. To this end, we used HemoLens to evaluate and delineate the phenotypical differences between WT and SCA mouse vessels. We further provide an example of HemoLens’ applicability in vascular research by presenting evidence of vasoconstriction and vasodilation of WT vessels in response to Phenylephrine and Acetylcholine, respectively. Using standard myography protocols and the RAMP system, we show that WT vessels are more compliant than SCA vessels in both static and dynamic conditions, which agrees with previously published work^42,73^. However, our isolated vessels did not reach critical intraluminal pressure to cause vessel rupture for true native burst pressure testing. This is likely attributed to the rupture pressure of mouse arteries being significantly higher than the 500 mmHg working range limit of the pressure sensors employed in our system^74^. Further, since most vascular tissue engineering is dedicated to creating biological alternatives from humans^75^, HemoLens was tuned to a frequency of 1Hz to closely mimic human in-vivo conditions^39^. If researchers aim to test the maximum burst pressure or replicate the frequency of murine heart rate of 10Hz^76^ of native or engineered vessels, HemoLens’ modularity allows users to upgrade the inline pressure sensors, upgrade RAMP’s motor control to provide faster rotation, and tune the Wheatstone bridge dynamic range to meet their needs.

Due to size limitations, genetic differences, and xenograft complications, tissue engineering is moving away from animal models to tissue-engineered vascular substitutes to study human-specific conditions, often incorporating human-derived cells and materials^7,9,77–79^. For this reason, we FRESH printed collagen-I vascular scaffolds as a proof-of-concept, small-diameter platform to study hypertensive disease progression. To compensate for the inherent mechanical weakness of collagen scaffolds, we designed our scaffolds to have a wall thickness larger than the reported wall thicknesses of native arteries, but still within ranges of recently published work on acellular vessels^80,81^. To direct our collagen scaffolds towards a more human hypertensive mechanical phenotype, we chemically crosslinked them with 4% PFA. This process resulted in a significantly higher burst pressure of 464 mmHg, nearly doubling that of our non-crosslinked scaffolds. Many chemical crosslinkers have been used to tune the mechanical properties of acellular scaffolds^82,83^. A 4% PFA solution was selected to emphasize two extreme mechanical phenotypes observed in blood vessels. These crosslinked acellular scaffolds were also less dynamically compliant than our non-crosslinked vessels, requiring a pathological pressure profile to distend similarly to the non-crosslinked vessels at a physiological pressure profile. The non-crosslinked collagen vessels tested were above physiological pressures but below the burst pressure required to achieve adequate safety factors for a human blood vessel equivalent^80^.

In addition to chemical crosslinking, cellularization of the non-crosslinked vessels should significantly improve the mechanical performance and shift the vessel phenotype toward more compliant native human vasculature. Collagen and elastin are two of the primary components that affect the elasticity of the vessel wall^80^. Previous reports have demonstrated that elastin deposition and increased fibril collagen within vascular grafts significantly shift their phenotype toward a more native human vasculature phenotype^84–87^. *In-vivo,* Elliott *et al.* showed that cellular ingrowth of their endothelialized fibrin vessel after a 24-week implantation had significantly increased the deposition of elastin, leading the remodeled conduit to have similar elasticity levels to that of native human aortas^84^. Similarly, *in-vitro*, Huang *et al.* demonstrated that cultured smooth muscle cells onto PGA scaffolds within a biaxial stretching bioreactor had increased fibrillar collagen production, which resulted in higher suture retention and distention rates similar to human vasculature^86^. The next phase of this research will focus on the generation of FRESH printed cellularized small-diameter vessels and the use of HemoLens as a mechanical characterization platform.

Parsing out mechanical differences in vessel wall behavior are critical for designing appropriate loading regimes for understanding aspects of vascular mechanobiology associated with disease progression across vascular beds. Analysis of material stiffness with advanced biomechanical analysis revealed our ability implement HemoLens to capture critical differences in non-linear vessel mechanics behavior that are key to understanding differences across vascular phenotypes. By comparing WT and SCA samples, we found that stiffening was exacerbated at the high pressures present in the diseased case. For bioprinted scaffolds, stress vs strain behavior revealed a behavior that is typically considered stiffer, through a leftward shift of the stress vs stretch curve. However, due to decreased deformation, similar to the decreased compliance observed in dynamic testing, the calculated values of material stiffness were lower with crosslinking. The leftward shift of the bioprinted scaffolds toward a collagen dominated region reflects the material composition of our printed scaffolds, which is only collagen-I. Using this data as a guide and FRESH bioprinting’s flexibility in printing various ECM bioink, we can tune the mechanics by adding additional ECM materials. Biomaterials such as collagen-II & III^88^, fibrin^89^, and elastin peptides^90^ can be directly printed via FRESH or supplemented in our bioink to shift our bioprinted scaffold mechanics toward a more physiological regime shown in our WT samples.

In this report, we present an open-source vascular characterization device called HemoLens, and provide several examples of feasibility and types of data derived using the HemoLens system. HemoLens is an inexpensive, comprehensive platform for the mechanical assessment of native and tissue engineered vasculature. Several relevant use cases of standard vascular characterization techniques were conducted on both native vessels and bioprintined scaffolds. While we focused on evaluation of small-diameter vessels in this work, the modular nature of HemoLens enables the characterization of larger-diameter scaffolds and blood vessels without modification. Moreover, although systems capable of delivering cyclic pressure waves for preconditioning and dynamic compliance testing have been developed previously^50,91^, to our knowledge, this is the first published report detailing the design, construction, and testing of a complete low-cost myography system that incorporates pulse pressure controlled via intraluminal flow. As a characterization platform, HemoLens significantly decreases the barrier of entry in vascular research by providing a rapidly deployable, low-cost myography system. As of now, HemoLens can only test a single vessel. However, multiple HemoLens systems can be built to increase experimentation throughput at a fraction of the cost of a single commercial or open-source system. Further, the innovative RAMP system provides HemoLens with additional functionality beyond commercial or open-source systems by producing cyclic pressure waves that more closely recapitulate the dynamic environment of the body. We hope researchers will capitalize on HemoLens to aid in developing TEBVs, validate hypertension vascular models, characterize cellularized grafts, or other myography-based applications.

## Materials and Methods

### Data Availability

All custom Python scripts and software used are available at our lab’s GitHub account post publication (https://github.com/STEL-Pitt-BioE/HemoLens). All data are available in the main text and the supplementary materials. STL files for 3D bioprinter hardware modifications, collagen constructs, and perfusion systems are available under an open-source CC-BY-SA license at Zenodo.com. Additional open-source 3D models can be found at www.ShiwarskiLab.com post publication.

### Device construction

HemoLens was ideated and modeled in Fusion360 (Autodesk). The primary structure of its components was 3D printed from Polylactic acid (PLA) or Polyethylene Terephthalate Glycol (PETG) with a BambuLab X1 3D printer. Commercial components, including screws, linear rails, and nuts, were added as needed. A build guide and STEP files for HemoLens’ are provided as supplementary materials and deposited in our GitHub repository (https://github.com/STEL-Pitt-BioE/HemoLens).

### Micromanipulator Control

Micromanipulator precision was measured as a function of linear movement repeatability in 1 mm and 0.25 mm increments, with an identical procedure employed for each of the micromanipulator’s three axes. HemoLens’ corner feet were magnetically attached to the base platform, and the system was placed flat on a laboratory bench. An erasable marker was used to demarcate one starting and three quarter-turn positions for the three knurl thumbs screws through which the X, Y, and Z bolts thread. A digital indicator (Mitutoyo model# S112S) was zeroed and placed perpendicular to the back of the manipulator on HemoLens’ vessel outlet side. The X-axis M6 bolt was manually turned to the first quarter-turn position, and the linear travel displayed on the digital indicator was recorded. This was repeated ten times; the digital indicator was reset to 0 mm after each recorded measurement. The X-axis bolt’s position was reset, and the experiment was repeated ten additional times with a full rotation. This procedure was replicated with the Y and Z-axis bolts, with the digital indicator placed on the corresponding side of the manipulator.

### Vessel imaging

Prior to each experiment, an M12 Lens 1/2.3” (ArduCam #LN024) was attached to a 12MP USB camera module (ArduCam #BO433), and the module was attached to HemoLens’ camera gantry. The camera gantry was magnetically attached to the camera stand, which was then magnetically attached to one side of HemoLens’ base platform. Vessels are properly illuminated from below with a USB powered LED Strip (Topai #COBW1-480-5000K) that activates when HemoLens’ is turned on. With the vessels mounted, Vasotracker software (V1.4.0) is started and OpenCV is used to establish the connection between the software and the camera system. Focus was established by moving the camera gantry up and down until a clear image of the vessel walls could be seen on the live image within Vasotracker. Vasotracker’s diameter-tracking algorithm was engaged, which plotted vessel ID and OD. The scale was adjusted in Vasotracker to corroborate previously measured nominal dimensions using a reference calibration slide.

### Temperature testing

Tubing lines were constructed from 1/16” silicone tubing and connected to each plastic perfusion port embedded in the bath insert. Each tubing line was then connected to two peristaltic pumps (Gikfun, GK1028). More tubing was then connected from each peristaltic pump to a single reservoir placed inside the water bath (IVYX Scientific). 30-gauge needles (JensonGlobal #JG30-0.5HPX) were press-fit into each HemoLens’ micromanipulator. A silicone test vessel was then press-fit and sutured onto the protruding side of the needles. Luer connectors, along with 1/16” silicone tubing, were then connected to the non-vessel side of the inlet and outlet needle and fed to an external small-volume waterproof aquarium pump (DOMICA #FR-PPU12-188). A 10 mL BD syringe was attached to the pump inlet via a custom 3D printed adapter, which the vessel outlet tubing was then attached via the Luer connection. Vessel inlet tubing was then connected to the pump outlet, which was retrofitted to house a female luer connector. The pump was placed in the water bath, primed with 5 mL 1x PBS (Accuris lot #2084806191), and engaged via HemoLens’ LCD to circulate fluid through the test vessel. The water bath was filled with water and set to a temperature of 47 °C. HemoLens’ bath insert and bath reservoir were both filled with 1x PBS, and the peristaltic pumps were engaged via HemoLens’ LCD to circulate fluid through the bath. Thermal probes were placed inside HemoLens’ bath, the bath reservoir, and the vessel line reservoir. Temperature was recorded every 5 minutes until bath and reservoir temperatures reached 37 °C.

### Vessel mounting

For experiments in both static and pulsatile pressure, the HemoLens bath was filled with 30 mL of 1x PBS. One 30-gauge dispensing needle was press-fit to each micromanipulator, and two 7-0 silk sutures (Braintree Scientific #SUT-S103). were lightly looped over each needle. Both native and FRESH printed vessels were gently placed into the HemoLens bath. Using clean forceps, each end of the vessel was slid over the 30-gauge needles. Using an overhead microscope, both ends of each vessel were manually sutured by double-knotting of the 7-0 silk suture After mounting, the bath was set to 47 °C, and pumps were engaged for at least 30 minutes to allow the bath to warm to physiological temperature. HemoLens’ camera gantry was attached to the camera stand, and the Vasotracker application was opened to engage the camera. Imaging settings were adjusted as previously described.

### Mouse vessel excision

Twenty-week-old C57BL/6J (stock no. 000664) mice were ordered from the Jackson Laboratory (Bar Harbor, ME). Sickle (SS) Townes knock-in mice were bred and maintained at the University of Pittsburgh. Bone marrow transplant was performed as previously described^92^. All animal experiments were reviewed and approved by the Institutional Animal Care and Use Committee at the University of Pittsburgh. The mice were all supplied a normal chow diet (no. 5234), had free access to drinking water. Mice were housed in pathogen free conditions in accordance with the Guide for the Care and Use of Laboratory Animals from the Department of Laboratory Animal Research at the University of Pittsburgh. The mice were euthanized, and both carotid arteries were excised. In our experiments, the wildtype arteries are normotensive, and arteries from sickle cell mice were considered hypertensive vessels. Each excised vessel was stored in 1x PBS at 4 °C. All vessels were stored in 1x PBS and used within 72 hours of excision. Vessels used for vasoreactivity were excised and used within 6 hours to minimize cellular function loss.

### Static Pressure Experiments

All reported experiments within this paper were conducted with 3 mL syringes (BD #309657). However, syringes (BD) up to 10 mL are compatible with the ratchet. Before vessel mounting, a 3 mL syringe (BD) was filled with 1x PBS and placed inside the ratchet system housing. The syringe was held in place via a designed magnetic clamp. A 3-way stopcock with 1/16” tubing was attached to the syringe and connected to a 30-gauge needle already attached to the micromanipulators. Air was purged from the tubing lines by compressing the syringe with the iPRS rachet system. The outflow tube was connected and purged of air in a similar manner. For vessel mounting, all tubing lines were sealed by setting each stopcock to the close position to ensure no air or water escaped during mounting. Vessels were mounted as described above. After the vessels were mounted, stopcocks were set to the open position, and the ratchet was continuously turned counterclockwise to plunge the syringe and perfuse the loaded 1x PBS through the vessel and into a collection beaker to remove any remaining air bubbles. The outlet line was then closed by turning the stopcock valve to the close position, ensuring a baseline pressure as close to 0 mmHg as possible. Systemic pressure was increased in discrete increments by rotating the knurled handled of the ratchet system counterclockwise. The pressure was held for 20 second intervals and increased until either burst pressure or systemic pressure of ∼500 mmHg was reached. Vessel diameters and systemic pressure were tracked using VasoTracker pressure myography software. This process was repeated for both native and FRESH printed samples. Software settings were adjusted to ensure the starting outer and inner diameters were aligned with the measured values of each carotid or printed vessel at atmospheric pressure. The maximum pressure before rupture was taken as the burst pressure.

### Vasoreactivity of normotensive vessels under static and dynamic conditions

To test the vasoreactivity of mouse carotid arteries, the mounted vessels were exposed to increasing concentrations of vasodilators and vasoconstrictors as previously described with slight modifications^45,46^. Normotensive carotid arteries were excised from c57bl/6J mice and placed in a physiological salt solution (PSS) composed of 118.4 mM NaCl (FisherSci #L23071), 6.0 mM d-glucose (Sigma #101979554), 4.0 mM NaHCO_3_ (Sigma #SLBM8267V), 4.7 mM KCL (Sigma # P4504), 1.2 mM MgSO_4_ (Sigma #M2773), 1.2 mM KH_2_PO_4_ (FisherSci #976446), and 2.0 mM CaCl_2_ (Sigma #MKBR9337V), buffered with 10 mM HEPES. pH of PSS was adjusted to 7.42 with 5N NaOH (FisherSci # 219185). Before adding the excised vessels into the bath, two 7-0 silk sutures were loosely tied to each 30-gauge needle protruding from the micromanipulators. The excised vessels were removed from their microtube and gently placed into the HemoLens bath containing 30 mL of PSS to float. Using clean forceps, each vessel was manually canulated to the 30-gauge needles and sutured tight to the needle. After mounting, the vessels were allowed to rest for 30 minutes to heat up to 37 °C in PSS. Once the physiological temperature was reached, the vessel pumps were turned on, and the systemic pressure was set to 84 mmHg. Normotensive vessels were exposed to 10^-7^ -10^-4^ M Phenylephrine (Sigma #SLBC3670V) in 2 minute intervals to induce maximum constriction before the introduction of vasodilation. After maximal vasoconstriction, vessels were given cumulatively increasing doses of Acetylcholine (Sigma #BCCD7863) 10^-7^ – 10^-4^ M in 2-minute intervals to determine maximum vasodilation within the working range. All vasoactive drugs were administered directly into the recirculating bath. For Phenylephrine, the data is reported as percentage constriction, with 100% being the highest degree of constriction and 0% as the lowest. For Acetylcholine, the data is reported as percentage relaxation of the vessel, with 0% reported as the starting diameter and 100% denoting the largest diameter. For all vasoreactivity studies vessels were recorded with the Windows 11 Camera App (Microsoft). FIJI (NIH) or Premiere Pro (Adobe) were used to convert recorded video into a usable format for Vasotracker’s offline analyzer.

### RAMP system

The RAMP system was engaged for experiments utilizing pulsatile pressure. After vessels were mounted and sutured, and HemoLens’ heating was prepared as previously described, pressure sensors (Honeywell #26PCDFG5G) were placed near the vessel inlet and outlet, connected to the 30-gauge needles via additional 1/16” tubing. HemoLens’ aquarium pump (DOMICA #AM-118) was primed with 5 mL PBS and activated via HemoLens’ LCD. Vessel outlet tubing was threaded through a designed circular clamp placed shortly after the vessel outlet. The clamp was manually cinched by tightening two M3 screws, increasing pressure to a baseline 80 mmHg as determined by continuous readings displayed on HemoLens’ LCD. The RAMP system was placed shortly before the clamp, and the vessel tubing was held directly under RAMP’s occlusion arm by two tubing holders designed into the base. The RAMP system was activated via HemoLens’ LCD , causing the RAMP’s arm to swing and pinch the outlet tubing housed in its slot. To verify the profile of RAMP’s pressure waves, pressure readings were recorded via a custom Python script. Unless otherwise stated, RAMP was tuned to mimic the healthy physiology of 80 mmHg diastolic and 120 mmHg systolic pressures were for all experiments.

### Dynamic compliance measurements

To perform dynamic compliance testing, vessels and HemoLens were prepared as outlined above. With the pumps on, baseline pressure is adjusted to roughly 80 mmHg using our designed small clamp to slightly occlude the outlet line until the desired baseline pressure is achieved. Baseline pressure is monitored on HemoLen’s LCD screen during adjustment. Once set, the HemoLens system is restarted, and the pump and RAMP system are activated during the normal HemoLens startup process. In our experiments, due to Vasotracker’s limitation in video frame rate, vessel distension was recorded for 1 minute with the Windows 11 camera app (Microsoft), and diameters were calculated using Vasotracker’s offline analyzer^24^. FIJI (NIH) or Premiere Pro (Adobe) were used to convert recorded video into a useable format for Vasotracker’s offline analyzer. Real time pressure during the experiment was recorded using a custom Python script. The outer diameter was used in all dynamic compliance calculations for both native and FRESH printed scaffolds as no significant differences were observed between inner and outer diameter measurements previously. Software settings in the offline analyzer were adjusted to ensure the starting diameters at baseline were consistent with results taken from burst pressure measurements. A custom Python script was used to filter and compile the recorded diameter changes and provide a list of the diameter changes of each peak. Diameter changes were averaged over the number of sampled peaks. Another Python script used fast Fourier transform (FFT) to calculate pulse frequency of vessel distention. Dynamic compliance (*Dc*) (mmHg^-1^) was calculated as previously described^50^.

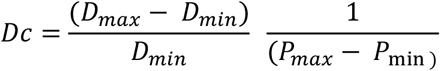

where D_max_ = the recorded diameter of each peak (µm), D_min_ = the baseline recorded diameter (µm), P_max_ = maximum pressure peak (mmHg), and P_min_ = minimum or baseline pressure (mmHg).

### Characterization of Mechanical Behavior

To demonstrate the usability of data gathered with the HemoLens platform in advanced analysis, we used a biomechanical framework to characterize the mechanical behavior of sample specimens from a variety of testing conditions: normotensive carotid, hypertensive carotid, printed vessel without crosslinking, and printed vessel with crosslinking. Our framework closely follows that developed previously for biomechanical phenotyping of murine blood vessels^66^. Loading protocols in this work were reduced, as our focus was on development and validation of the HemoLens device rather than on comprehensive testing procedures. Therefore, pressure and loaded outer diameter from a single testing cycle of each specimen were used for fitting with a constant axial stretch. A four fiber family-based (4FF) stored energy densify function, extensively used to simulate mechanical behavior of vascular tissue, was used to capture the measured experiments from this work. We fit the material parameters of the 4FF model by matching the model-generated pressure vs diameter behavior to that gathered experimentally with numerical optimization. Linearized stiffness, which is a measure of the material stiffness in the loaded configuration, was calculated according to prior methods at different pressure states as the main output for comparison^66,93^. This material stiffness was calculated at different pressures, which is necessary for non-linear materials to observe differences in behavior across loading states.

### Gelatin microparticle support generation for FRESH bioprinting

FRESH v2.0 support bath was prepared with a coacervation method as previously described with slight modifications^59^. A liter (1000 mL) of support solution was made by dissolving 3% (w/v) Gelatin (Fisher Sci #235321), 0.3% (w/v) Gum Arabica (Sigma #102530584), and 0.125% (w/v) Pluronic F127 (Sigma #102525773) in warmed 50%(v/v) Ethanol-Milli-Q water solution. The solution’s pH was adjusted to 5.35 with 5 N hydrochloric acid (FischerChemical lot# 226234). An overhead stirrer (IKA) was used to mix the solution at 580 RPM overnight at room temperature. Polyvinylidene chloride (PDVC) wrap was used to seal the solution to minimize evaporation during overnight mixing. The support was transferred into four 250 ml containers and spun at 500 x g for 3 minutes to pellet the gelatin support. The supernatant was decanted from the containers. The gelatin support was washed three more times by filling each container with 125 mL of Milli-Q water and shaking vigorously by hand before being spun down at 750 x g for 2 minutes. The supernatant was removed after each spin. Compacted gelatin support was resuspended in a 1% Penn Strep (Gibco # 208330) Milli-Q water solution and held at 4 °C until needed. Prior to printing, the uncompacted gelatin support was compacted by centrifugation at 1000xg for 1 minute, and the supernatant was decanted. The compacted support was resuspended in 50mM HEPES (Sigma #1003551688) and degassed in a vacuum chamber for 30 minutes, followed by centrifugation at 1850 x g for 3 minutes. After the final centrifugation step, the supernatant was removed, and the compacted support was transferred into a 35 mm printing dish.

### Collagen bioink preparation

Collagen-I bioink was purchased as LifeInk 260 (Advanced Biomatrix #5358). No modification or dilution was done to the bioink, and it was used as packaged by the manufacturer. The bioink was centrifuged for 5 minutes at 3000 g to ensure the removal of bubbles from the solution. The collagen-I bioink was transferred to a 1mL gastight glass syringe (Hamilton #1089113) for printing.

### FRESH bioprinting of collagen vessels

Test vessels were designed in an array fashion nested within a support frame using Fusion360 (AutoCAD) with 450 µm inner diameter and 1200 µm outer diameter. Machine pathing was visualized and Gcode Cura v5.7 (Ultimaker). For printing, Gcode was uploaded to our custom-built bioprinter via a universal serial bus (USB) flash drive. Once the print was complete, the collagen vessels were allowed to rest at room temperature for at least 30 minutes to continue crosslinking. After room temperature incubation, the prints were transferred to a benchtop incubator set at 37 °C to melt away the gelatin support. After 1 hour of incubation at 37 °C, gelatin support was washed away with repeated 50% 50 mM HEPES (Sigma #1003551688) exchanges every 30-45 minutes thereafter until no more gelatin was apparent in the HEPES bath. Once cleared of gelatin support, printed scaffolds were stored in fresh 50 mM HEPES at 4 °C until needed.

### Imaging and measuring of printed vessels

Before imaging, prints were washed one final time in fresh 50 mM HEPES on a rocker at 37 °C to ensure the removal of any remaining gelatin microparticles within the lumen of the prints. A 100% 50 mM HEPES exchange was conducted prior to imaging. Printed vessels were imaged using a brightfield stereomicroscope (Leica M205 FA with a 1X objective and K8 camera). Print quality was assessed based on the integrity of the printed scaffold’s walls. Any holes, voids, or separation of layers of the wall were considered to be a failed print. The inner and outer diameters of the printed vessels were measured using the line profile measurement tool in LasX v3.9 (Leica) software. A total of 5 measurements across the length of scaffold was used to generate average diameters and standard deviations.

### Paraformaldehyde fixing of FRESH printed collagen scaffolds

A fixing solution of 4% Paraformaldehyde was created by diluting a 16% Paraformaldehyde (ThermoScientific lot# T15K021) stock concentration with MiliQ water. The fixing solution was stored at room temperature until needed. Printed scaffolds were immersed in 4% PFA for 30 minutes at room temperature. Printed scaffolds were washed 3x in 50 mM HEPES for 2-3 minutes each and stored in fresh 50 mM HEPES at 4 °C until needed.

### Data Analysis

Custom Python scripts were used to calculate average diameter changes and produce the frequency spectra via fast Fourier transforms (FFT). Please see data availability section for details to accessing custom Python scripts. Statistical and graphical analyses were conducted using Prism v10 (GraphPad) and Excel v16 (Microsoft) software. Statistical tests were chosen based on sample size, normality of the data set, and data requirements. For pressure increases in syringe diameter, a student’s T-test was used to compare pressure increases between 3 mL and 1 mL syringe types. Significance was determined as p<0.05. For average diameter and dynamic compliance, our sample size of pulse peaks (technical replicates) was above n=30, sufficient for the central limit theorem (CLT) to assume our data set reaches a standard Gaussian distribution. A one-way analysis of variance (ANOVA) with Tukey’s multiple comparisons was used to determine the significance of average diameter and dynamic compliance. Significance was defined as p<0.05 for all comparisons.

## Acknowledgments

We thank members of the Shiwarski lab for their input, support, and guidance during the completion of this work and preparation of the manuscript.

This works was supported by the National Heart, Lung, And Blood Institute of the National Institutes of Health (Award Numbers R00HL155777 (D.J.S), 5T32HL149648-05 (B.D.C), R35 HL161177 (A.C.S), and the Department of Defense (DoD) under award number HQ00342110020 (A.J.P) issued by the Office of Naval Research.

## Author Contributions

Conceptualization, A.P.T, L.A.M, F.A, B.D.C, and D.J.S; Methodology: A.P.T, L.A.M, and D.J.S Software: A.P.T, J.M.S, and L.A.M; Formal Analysis: A.J.P, D.J.S; Investigation A.P.T, L.A.M, and S.H; Resources: S.H, A.C.S, and D.J.S; Data curation: A.P.T and D.J.S; Writing-Original Draft: A.P.T, L.A.M, J.M.S, S.H, A.C.S, and D.J.S; Visualization: A.P.T, L.A.M, J.M.S, and D.J.S; Funding Acquisition: A.C.S and D.J.S; Supervision: A.C.S and D.J.S.

## REFERENCES

1. Staessen, J.A., Wang, J., Bianchi, G., and Birkenhager, W.H. (2003). Essential hypertension. Lancet 361, 1629–1641. 10.1016/S0140-6736(03)13302-8.

2. Brown, I.A.M., Diederich, L., Good, M.E., DeLalio, L.J., Murphy, S.A., Cortese-Krott, M.M., Hall, J.L., Le, T.H., and Isakson, B.E. (2018). Vascular Smooth Muscle Remodeling in Conductive and Resistance Arteries in Hypertension. Arterioscler Thromb Vasc Biol 38, 1969–1985. 10.1161/ATVBAHA.118.311229.

3. Camasao, D.B., and Mantovani, D. (2021). The mechanical characterization of blood vessels and their substitutes in the continuous quest for physiological-relevant performances. A critical review. Mater Today Bio 10, 100106. 10.1016/j.mtbio.2021.100106.

4. Humphrey, J.D. (2021). Mechanisms of Vascular Remodeling in Hypertension. Am J Hypertens 34, 432–441. 10.1093/ajh/hpaa195.

5. Xie, S.A., Zhang, T., Wang, J., Zhao, F., Zhang, Y.P., Yao, W.J., Hur, S.S., Yeh, Y.T., Pang, W., Zheng, L.S., et al. (2018). Matrix stiffness determines the phenotype of vascular smooth muscle cell in vitro and in vivo: Role of DNA methyltransferase 1. Biomaterials 155, 203–216. 10.1016/j.biomaterials.2017.11.033.

6. Savoia, C., Sada, L., Zezza, L., Pucci, L., Lauri, F.M., Befani, A., Alonzo, A., and Volpe, M. (2011). Vascular inflammation and endothelial dysfunction in experimental hypertension. Int J Hypertens 2011, 281240. 10.4061/2011/281240.

7. Williams, S.M., Haines, J.L., and Moore, J.H. (2004). The use of animal models in the study of complex disease: all else is never equal or why do so many human studies fail to replicate animal findings? Bioessays 26, 170–179. 10.1002/bies.10401.

8. Monassier, L., Combe, R., and Fertak, L.E. (2006). Mouse models of hypertension. Drug Discovery Today: Disease Models 3, 273–281. 10.1016/j.ddmod.2006.10.008.

9. Seok, J., Warren, H.S., Cuenca, A.G., Mindrinos, M.N., Baker, H.V., Xu, W., Richards, D.R., McDonald-Smith, G.P., Gao, H., Hennessy, L., et al. (2013). Genomic responses in mouse models poorly mimic human inflammatory diseases. Proc Natl Acad Sci U S A 110, 3507–3512. 10.1073/pnas.1222878110.

10. Caddeo, S., Boffito, M., and Sartori, S. (2017). Tissue Engineering Approaches in the Design of Healthy and Pathological In Vitro Tissue Models. Front Bioeng Biotechnol 5, 40. 10.3389/fbioe.2017.00040.

11. Joyce, K., Buljovcic, Z., Rosic, G., Kaszkin-Bettag, M., and Pandit, A. (2023). Issues with Tissues: Trends in Tissue-Engineered Products in Clinical Trials in the European Union. Tissue Eng Part B Rev 29, 78–88. 10.1089/ten.TEB.2022.0094.

12. Niklason, L.E., and Lawson, J.H. (2020). Bioengineered human blood vessels. Science 370. 10.1126/science.aaw8682.

13. Jia, W., Gungor-Ozkerim, P.S., Zhang, Y.S., Yue, K., Zhu, K., Liu, W., Pi, Q., Byambaa, B., Dokmeci, M.R., Shin, S.R., and Khademhosseini, A. (2016). Direct 3D bioprinting of perfusable vascular constructs using a blend bioink. Biomaterials 106, 58–68. 10.1016/j.biomaterials.2016.07.038.

14. Sundaram, S., Lee, J.H., Bjorge, I.M., Michas, C., Kim, S., Lammers, A., Mano, J.F., Eyckmans, J., White, A.E., and Chen, C.S. (2024). Sacrificial capillary pumps to engineer multiscalar biological forms. Nature 636, 361–367. 10.1038/s41586-024-08175-5.

15. Daniel J. Shiwarski, A.R.H., Joshua W. Tashman, Ezgi Bakirci, Samuel Moss, Brian D. Coffin, Adam W. Feinberg (2025). 3D bioprinting of collagen-based high-resolution internally perfusable scaffolds for engineering fully biologic tissue systems. Science Advances 11, 1–20. 10.1126/sciadv.adu5905.

16. Bagrat Grigoryan, S.J.P., Daniel C. Corbett, Daniel W. Sazer, Chelsea L. Fortin, Alexander J. Zaita1, Paul T. Greenfield1, Nicholas J. Calafat, John P. Gounley, Anderson H. Ta, Fredrik Johansson, Amanda Randles, Jessica E. Rosenkrantz, Jesse D. Louis-Rosenberg, Peter A. Galie, Kelly R. Stevens2 Jordan S. Miller (2019). Multivascular Networks and Functional Intravascular Topologies within Biocompatible Hydrogels. Science 364, 458–464.

17. Budoff, M.J., Alpert, B., Chirinos, J.A., Fernhall, B., Hamburg, N., Kario, K., Kullo, I., Matsushita, K., Miyoshi, T., Tanaka, H., et al. (2022). Clinical Applications Measuring Arterial Stiffness: An Expert Consensus for the Application of Cardio-Ankle Vascular Index. Am J Hypertens 35, 441–453. 10.1093/ajh/hpab178.

18. Bailey, A.L., and Smyth, S.S. (2012). Invasive coronary vasoreactivity testing to diagnose microvascular dysfunction in women. JACC Cardiovasc Interv 5, 654–655. 10.1016/j.jcin.2012.03.014.

19. Gleason, R.L., Gray, S.P., Wilson, E., and Humphrey, J.D. (2004). A multiaxial computer-controlled organ culture and biomechanical device for mouse carotid arteries. J Biomech Eng 126, 787–795. 10.1115/1.1824130.

20. Valsecchi, E., Biagiotti, M., Alessandrino, A., Gastaldi, D., Vena, P., and Freddi, G. (2022). Silk Vascular Grafts with Optimized Mechanical Properties for the Repair and Regeneration of Small Caliber Blood Vessels. Materials (Basel) 15. 10.3390/ma15103735.

21. Xu, L., Varkey, M., Jorgensen, A., Ju, J., Jin, Q., Park, J.H., Fu, Y., Zhang, G., Ke, D., Zhao, W., et al. (2020). Bioprinting small diameter blood vessel constructs with an endothelial and smooth muscle cell bilayer in a single step. Biofabrication 12, 045012. 10.1088/1758-5090/aba2b6.

22. Stoiber, M., Messner, B., Grasl, C., Gschlad, V., Bergmeister, H., Bernhard, D., and Schima, H. (2015). A method for mechanical characterization of small blood vessels and vascular grafts. Experimental Mechanics 55, 1591–1595. 10.1007/s11340-015-0053-x.

23. Jadeja, R.N., Rachakonda, V., Bagi, Z., and Khurana, S. (2015). Assessing Myogenic Response and Vasoactivity In Resistance Mesenteric Arteries Using Pressure Myography. J Vis Exp, e50997. 10.3791/50997.

24. Lawton, P.F., Lee, M.D., Saunter, C.D., Girkin, J.M., McCarron, J.G., and Wilson, C. (2019). VasoTracker, a Low-Cost and Open Source Pressure Myograph System for Vascular Physiology. Front Physiol 10, 99. 10.3389/fphys.2019.00099.

25. Wenceslau, C.F., McCarthy, C.G., Earley, S., England, S.K., Filosa, J.A., Goulopoulou, S., Gutterman, D.D., Isakson, B.E., Kanagy, N.L., Martinez-Lemus, L.A., et al. (2021). Guidelines for the measurement of vascular function and structure in isolated arteries and veins. American Journal of Physiology-Heart and Circulatory Physiology 321, H77–H111. 10.1152/ajpheart.01021.2020.

26. Olav L. Schjørring, R.C., and Ulf Simonsen (2015). Pressure Myography to Study the Function and Structure of Isolated Small Arteries. In Methods in Molecular Biology, V. Andrés, and B. Dorado, eds. (Humana Press), pp. 277–295.

27. Tashman, J.W., Shiwarski, D.J., and Feinberg, A.W. (2022). Development of a high-performance open-source 3D bioprinter. Sci Rep 12, 22652. 10.1038/s41598-022-26809-4.

28. Pearce, J.M. (2012). Building Research Equipment with Free, Open-Source Hardware. Science 337, 1303–1304. 10.1126/science.122818.

29. Stan Z. Li, and Jain, A.K. (2009). Blood Vessel Wall. In Encyclopedia of Biometrics, Stan Z. Li, ed. (Springer). 10.1007/978-0-387-73003-5_136.

30. Majesky, M.W., Dong, X.R., Hoglund, V., Mahoney, W.M., Jr., and Daum, G. (2011). The adventitia: a dynamic interface containing resident progenitor cells. Arterioscler Thromb Vasc Biol 31, 1530–1539. 10.1161/ATVBAHA.110.221549.

31. Seidel, C.L. (1997). Cellular Heterogeneity of the Vascular Tunica Media : Implications for Vessel Wall Repair. Arterioscler Thromb Vasc Biol 17, 1868–1871. 10.1161/01.ATV.17.10.1868.

32. Syedain, Z.H., Prunty, A., Li, J., and Tranquillo, R.T. (2021). Evaluation of the probe burst test as a measure of strength for a biologically-engineered vascular graft. J Mech Behav Biomed Mater 119, 104527. 10.1016/j.jmbbm.2021.104527.

33. Doppegieter, M., van Leeuwen, T.G., Aalders, M.C.G., de Vos, J., van Bavel, E.T., and Bakker, E. (2024). The impact of temperature on vascular function in connection with vascular laser treatment. Lasers Med Sci 39, 122. 10.1007/s10103-024-04070-7.

34. He, Q., Zhu, L., Lemons, D.E., and Weinbaum, S. (2002). Experimental measurements of the temperature variation along artery-vein pairs from 200 to 1000 microns diameter in rat hind limb. J Biomech Eng 124, 656–661. 10.1115/1.1517061.

35. Brueggemann, L.I., Mani, B.K., Haick, J., and Byron, K.L. (2012). Exploring arterial smooth muscle Kv7 potassium channel function using patch clamp electrophysiology and pressure myography. J Vis Exp, e4263. 10.3791/4263.

36. Seifu, D.G., Purnama, A., Mequanint, K., and Mantovani, D. (2013). Small-diameter vascular tissue engineering. Nat Rev Cardiol 10, 410–421. 10.1038/nrcardio.2013.77.

37. Bartolf-Kopp, M., and Jungst, T. (2024). The Past, Present, and Future of Tubular Melt Electrowritten Constructs to Mimic Small Diameter Blood Vessels - A Stable Process? Adv Healthc Mater 13, e2400426. 10.1002/adhm.202400426.

38. Bhandari, A.S.A.M.P. (2023). Vital Sign Assessment. NCBI Book Shelf, 1–10.

39. Herbert J. Levine, M., FACC (1997). Rest Heart Rate and Life Expectancy. Journal of the American College of Cardiology 30, 1104–1106. 10.1016/S0735-1097(97)00246-5.

40. Charlton, P.H., Mariscal Harana, J., Vennin, S., Li, Y., Chowienczyk, P., and Alastruey, J. (2019). Modeling arterial pulse waves in healthy aging: a database for in silico evaluation of hemodynamics and pulse wave indexes. Am J Physiol Heart Circ Physiol 317, H1062–H1085. 10.1152/ajpheart.00218.2019.

41. Kim, H.L. (2023). Arterial stiffness and hypertension. Clin Hypertens 29, 31. 10.1186/s40885-023-00258-1.

42. Hassona, M.D., Abouelnaga, Z.A., Elnakish, M.T., Awad, M.M., Alhaj, M., Goldschmidt-Clermont, P.J., and Hassanain, H. (2010). Vascular hypertrophy-associated hypertension of profilin1 transgenic mouse model leads to functional remodeling of peripheral arteries. Am J Physiol Heart Circ Physiol 298, H2112–2120. 10.1152/ajpheart.00016.2010.

43. Hannah Song, P.M.K., Suhaas Anbazhakan, Christian P. Rivera, Yundi Feng, Victor O. Omojola, Alexus A. Clark, Shuangyi Cai, Jada Selma, Rudolph L. Gleason Jr, Edward A. Botchwey, Yunlong Huo, Wenchang Tan, and Manu O. Platt (2020). Sickle Cell Anemia Mediates Carotid Artery Expansive Remodeling That Can Be Prevented by Inhibition of JNK (c-Jun N-Terminal Kinase). Arterioscler Thromb Vasc Biol 40, 1220–1230. 10.1161/ATVBAHA.120.314045.

44. Belizna, C., Loufrani, L., Ghali, A., Lahary, A., Primard, E., Louvel, J.P., Henrion, D., Levesque, H., and Ifrah, N. (2012). Arterial stiffness and stroke in sickle cell disease. Stroke 43, 1129–1130. 10.1161/STROKEAHA.111.635383.

45. Durgin, B.G., Hahn, S.A., Schmidt, H.M., Miller, M.P., Hafeez, N., Mathar, I., Freitag, D., Sandner, P., and Straub, A.C. (2019). Loss of smooth muscle CYB5R3 amplifies angiotensin II-induced hypertension by increasing sGC heme oxidation. JCI Insight 4. 10.1172/jci.insight.129183.

46. Durgin, B.G., Wood, K.C., Hahn, S.A., McMahon, B., Baust, J.J., and Straub, A.C. (2022). Smooth muscle cell CYB5R3 preserves cardiac and vascular function under chronic hypoxic stress. J Mol Cell Cardiol 162, 72–80. 10.1016/j.yjmcc.2021.09.005.

47. Sena, C.M., Goncalves, L., and Seica, R. (2022). Methods to evaluate vascular function: a crucial approach towards predictive, preventive, and personalised medicine. EPMA J 13, 209–235. 10.1007/s13167-022-00280-7.

48. Ku, D.N. (1997). Blood Flow in Arteries. Annual Review of Fluid Mechanics, 339–434. 10.1146/annurev.fluid.29.1.399.

49. Bernhard Sebastian, P.S.D. (2018). Microfluidics to Mimic Blood Flow in Health and Disease. Annual Review of Fluid Mechanics 50, 483–504. 10.1146/annurev-fluid-.

50. Soletti, L., Hong, Y., Guan, J., Stankus, J.J., El-Kurdi, M.S., Wagner, W.R., and Vorp, D.A. (2010). A bilayered elastomeric scaffold for tissue engineering of small diameter vascular grafts. Acta Biomater 6, 110–122. 10.1016/j.actbio.2009.06.026.

51. Ding, Z., Tang, N., Huang, J., Cao, X., and Wu, S. (2023). Global hotspots and emerging trends in 3D bioprinting research. Front Bioeng Biotechnol 11, 1169893. 10.3389/fbioe.2023.1169893.

52. Santoni, S., Gugliandolo, S.G., Sponchioni, M., Moscatelli, D., and Colosimo, B.M. (2021). 3D bioprinting: current status and trends—a guide to the literature and industrial practice. Bio-Design and Manufacturing 5, 14–42. 10.1007/s42242-021-00165-0.

53. Hudson, A.R., Shiwarski, D.J., Kramer, A.J., and Feinberg, A.W. (2025). Enhancing Viability in Static and Perfused 3D Tissue Constructs Using Sacrificial Gelatin Microparticles. ACS Biomater Sci Eng. 10.1021/acsbiomaterials.4c02169.

54. Christopher S. O’Bryan, T.B., Samuel Hart, Christopher P. Kabb, Kyle D. Schulze, Indrasena Chilakala, Brent S. Sumerlin, W. Gregory Sawyer, Thomas E. Angelini (2017). Self-assembled micro-organogels for 3D printing silicone structures. Science Advances 3, 1–8. 10.1126/sciadv.1602800.

55. Lin, N.Y.C., Homan, K.A., Robinson, S.S., Kolesky, D.B., Duarte, N., Moisan, A., and Lewis, J.A. (2019). Renal reabsorption in 3D vascularized proximal tubule models. Proc Natl Acad Sci U S A 116, 5399–5404. 10.1073/pnas.1815208116.

56. Mark A. Skylar-Scott, S.G.M.U., Lucy L. Nam, John H. Ahrens, Ryan L. Truby, Sarita Damaraju, Jennifer A. Lewis (2019). Biomanufacturing of organ-specific tissues with high cellular density and embedded vascular channels. Science Advances., 13. 10.1126/sciadv.aaw2459.

57. A. Lee, A.R.H., D. J. Shiwarski, J. W. Tashman, T. J. Hinton, S. Yerneni, J. M. Bliley, P. G. Campbell, A. W. Feinberg (2019). 3D bioprinting of collagen to rebuild components of the human heart. Science 365, 482–487. 10.1126/science.aav9051.

58. Thomas J. Hinton, Q.J., Rachelle N. Palchesko, Joon Hyung Park, Martin S. Grodzicki, Hao-Jan Shue, Mohamed H. Ramadan, Andrew R. Hudson, and Adam W. Feinberg (2015). Three-dimensional printing of complex biological structures by freeform reversible embedding of suspended hydrogels. Science Advances 1, 1–16. 10.1126/sciadv.1500758.

59. Shiwarski, D.J., Hudson, A.R., Tashman, J.W., and Feinberg, A.W. (2021). Emergence of FRESH 3D printing as a platform for advanced tissue biofabrication. APL Bioeng 5, 010904. 10.1063/5.0032777.

60. Wang, Y., Wang, Z., and Dong, Y. (2023). Collagen-Based Biomaterials for Tissue Engineering. ACS Biomater Sci Eng 9, 1132–1150. 10.1021/acsbiomaterials.2c00730.

61. Rico-Llanos, G.A., Borrego-González, S., Moncayo-Donoso, M., Becerra, J., and Visser, R. (2021). Collagen Type I Biomaterials as Scaffolds for Bone Tissue Engineering. Polymers 13. 10.3390/polym13040599.

62. Gurumurthy, B., and Janorkar, A.V. (2021). Improvements in mechanical properties of collagen-based scaffolds for tissue engineering. Current Opinion in Biomedical Engineering 17. 10.1016/j.cobme.2020.100253.

63. Wagenseil, J.E., and Mecham, R.P. (2009). Vascular extracellular matrix and arterial mechanics. Physiol Rev 89, 957–989. 10.1152/physrev.00041.2008.

64. Humphrey, J.D. (2003). Continuum biomechanics of soft biological tissues. Proceedings of the Royal Society of London. Series A: Mathematical, Physical and Engineering Sciences 459, 3–46. 10.1098/rspa.2002.1060.

65. Vatner, S.F., Zhang, J., Vyzas, C., Mishra, K., Graham, R.M., and Vatner, D.E. (2021). Vascular Stiffness in Aging and Disease. Front Physiol 12, 762437. 10.3389/fphys.2021.762437.

66. Ferruzzi, J., Bersi, M.R., and Humphrey, J.D. (2013). Biomechanical phenotyping of central arteries in health and disease: advantages of and methods for murine models. Ann Biomed Eng 41, 1311–1330. 10.1007/s10439-013-0799-1.

67. Martinez-Lemus, L.A. (2012). The dynamic structure of arterioles. Basic Clin Pharmacol Toxicol 110, 5–11. 10.1111/j.1742-7843.2011.00813.x.

68. Gonzalez-Fernandez, J.M. (1994). On the origin and dynamics of the vasomotion of small arteries. Mathematical Biosciences 119, 127–167. 10.1016/0025-5564(94)90074-4.

69. Jiang, Z., Diggle, B., Tan, M.L., Viktorova, J., Bennett, C.W., and Connal, L.A. (2020). Extrusion 3D Printing of Polymeric Materials with Advanced Properties. Adv Sci (Weinh) 7, 2001379. 10.1002/advs.202001379.

70. Kasmi, S., Ginoux, G., Allaoui, S., and Alix, S. (2021). Investigation of 3D printing strategy on the mechanical performance of coextruded continuous carbon fiber reinforced PETG. Journal of Applied Polymer Science 138. 10.1002/app.50955.

71. Vidakis, N., Petousis, M., and Kechagias, J.D. (2022). A comprehensive investigation of the 3D printing parameters’ effects on the mechanical response of polycarbonate in fused filament fabrication. Progress in Additive Manufacturing 7, 713–722. 10.1007/s40964-021-00258-3.

72. Yang, G., Mahadik, B., Choi, J.Y., and Fisher, J.P. (2020). Vascularization in tissue engineering: fundamentals and state-of-art. Prog Biomed Eng (Bristol) 2. 10.1088/2516-1091/ab5637.

73. Steppan, J., Jandu, S., Savage, W., Wang, H., Kang, S., Narayanan, R., Nyhan, D., and Santhanam, L. (2020). Restoring Blood Pressure in Hypertensive Mice Fails to Fully Reverse Vascular Stiffness. Front Physiol 11, 824. 10.3389/fphys.2020.00824.

74. Lane, B.A., Uline, M.J., Wang, X., Shazly, T., Vyavahare, N.R., and Eberth, J.F. (2021). The Association Between Curvature and Rupture in a Murine Model of Abdominal Aortic Aneurysm and Dissection. Exp Mech 61, 203–216. 10.1007/s11340-020-00661-x.

75. Fleischer, S., Tavakol, D.N., and Vunjak-Novakovic, G. (2020). From arteries to capillaries: approaches to engineering human vasculature. Adv Funct Mater 30. 10.1002/adfm.201910811.

76. Janssen, P.M., Biesiadecki, B.J., Ziolo, M.T., and Davis, J.P. (2016). The Need for Speed: Mice, Men, and Myocardial Kinetic Reserve. Circ Res 119, 418–421. 10.1161/CIRCRESAHA.116.309126.

77. Naegeli, K.M., Kural, M.H., Li, Y., Wang, J., Hugentobler, E.A., and Niklason, L.E. (2022). Bioengineering Human Tissues and the Future of Vascular Replacement. Circ Res 131, 109–126. 10.1161/CIRCRESAHA.121.319984.

78. Byrom, M.J., Bannon, P.G., White, G.H., and Ng, M.K. (2010). Animal models for the assessment of novel vascular conduits. J Vasc Surg 52, 176–195. 10.1016/j.jvs.2009.10.080.

79. Fishman, J.A., Scobie, L., and Takeuchi, Y. (2012). Xenotransplantation-associated infectious risk: a WHO consultation. Xenotransplantation 19, 72–81. 10.1111/j.1399-3089.2012.00693.x.

80. Latimer, C.A., Nelson, M., Moore, C.M., and Martin, K.E. (2014). Effect of collagen and elastin content on the burst pressure of human blood vessel seals formed with a bipolar tissue sealing system. J Surg Res 186, 73–80. 10.1016/j.jss.2013.08.003.

81. Regenberg, M.C., Wilhelmi, M., Hilfiker, A., Haverich, A., and Aper, T. (2023). Development, comparative structural analysis, and first in vivo evaluation of acellular implanted highly compacted fibrin tubes for arterial bypass grafting. J Mech Behav Biomed Mater 148, 106199. 10.1016/j.jmbbm.2023.106199.

82. Bax, D.V., Davidenko, N., Hamaia, S.W., Farndale, R.W., Best, S.M., and Cameron, R.E. (2019). Impact of UV- and carbodiimide-based crosslinking on the integrin-binding properties of collagen-based materials. Acta Biomater 100, 280–291. 10.1016/j.actbio.2019.09.046.

83. Davidenko, N., Schuster, C.F., Bax, D.V., Raynal, N., Farndale, R.W., Best, S.M., and Cameron, R.E. (2015). Control of crosslinking for tailoring collagen-based scaffolds stability and mechanics. Acta Biomater 25, 131–142. 10.1016/j.actbio.2015.07.034.

84. Morgan B. Elliott, B.G., Takuma Fukunishi, Djahida Bedja, Abhilash Suresh, Theresa Chen, Takahiro Inoue, Harry C. Dietz, Lakshmi Santhanam, Hai-Quan Mao, Narutoshi Hibino, and Sharon Gerecht (2022). Tissue engineered vascular grafts transform into autologous neovessels capable of native function and growth. Communications Medicine 2. 10.1038/s43856-021-00063-7.

85. Venkataraman, L., Bashur, C.A., and Ramamurthi, A. (2014). Impact of cyclic stretch on induced elastogenesis within collagenous conduits. Tissue Eng Part A 20, 1403–1415. 10.1089/ten.TEA.2013.0294.

86. Huang, A.H., Balestrini, J.L., Udelsman, B.V., Zhou, K.C., Zhao, L., Ferruzzi, J., Starcher, B.C., Levene, M.J., Humphrey, J.D., and Niklason, L.E. (2016). Biaxial Stretch Improves Elastic Fiber Maturation, Collagen Arrangement, and Mechanical Properties in Engineered Arteries. Tissue Eng Part C Methods 22, 524–533. 10.1089/ten.TEC.2015.0309.

87. Simon P. Hoerstrupa, G.Z.È., Ralf Sodian, Andrea M. Schnellc, Ju Èrg Gru Ènenfeldera, Marko I. Turina (2001). Tissue engineering of small caliber vascular grafts. Eur J Cardiothorac Surg 20, 164–169.

88. Moss, S.P., Shiwarski, D.J., and Feinberg, A.W. (2024). FRESH 3D Bioprinting of Collagen Types I, II, and III. ACS Biomater Sci Eng. 10.1021/acsbiomaterials.4c01826.

89. Freeman, S., Ramos, R., Alexis Chando, P., Zhou, L., Reeser, K., Jin, S., Soman, P., and Ye, K. (2019). A bioink blend for rotary 3D bioprinting tissue engineered small-diameter vascular constructs. Acta Biomater 95, 152–164. 10.1016/j.actbio.2019.06.052.

90. Cai, L., Dinh, C.B., and Heilshorn, S.C. (2014). One-pot Synthesis of Elastin-like Polypeptide Hydrogels with Grafted VEGF-Mimetic Peptides. Biomater Sci 2, 757–765. 10.1039/C3BM60293A.

91. Roger N. Baird, I.G.K., Gilbert J. L’Italien, and William M. Abott (1977). Dynamic compliance of arterial grafts. American journal of physiology 233, H568–H572. 10.1152/ajpheart.1977.233.5.H568.

92. Schmidt, H.M., DeVallance, E.R., Lewis, S.E., Wood, K.C., Annarapu, G.K., Carreno, M., Hahn, S.A., Seman, M., Maxwell, B.A., Hileman, E.A., et al. (2023). Release of hepatic xanthine oxidase (XO) to the circulation is protective in intravascular hemolytic crisis. Redox Biol 62, 102636. 10.1016/j.redox.2023.102636.

93. Baek, S., Gleason, R.L., Rajagopal, K.R., and Humphrey, J.D. (2007). Theory of small on large: Potential utility in computations of fluid–solid interactions in arteries. Computer Methods in Applied Mechanics and Engineering 196, 3070–3078. 10.1016/j.cma.2006.06.018.

