## Supplemental Figure and Legends for "Development of an Open-source Low-cost Pressure Myography and Cardiac Flow Simulator, HemoLens, for Mechanical Characterization of Native and Engineered Blood Vessels"

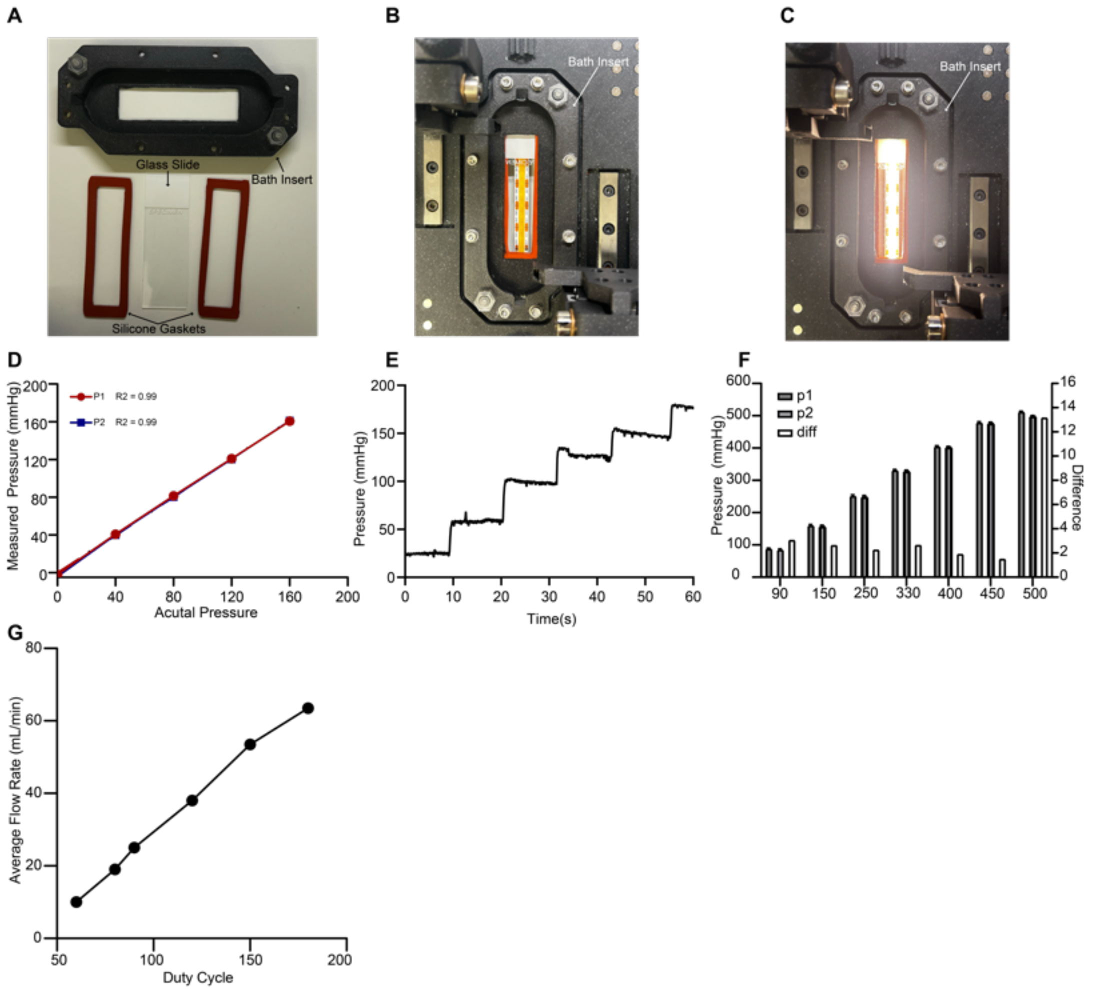


**S1 | A)** HemoLens' 3D printed bath insert, including glass slide and rubber gasket. **B)** HemoLen's bath insert is assembled into the 3D printed HemoLens base that is filled with liquid. **C)**. Light passing through liquid filled HemoLens' bath insert. **D)** The calibration curve of inline pressure sensors (p1 and p2) shows $r^2$ values of both > 0.98. **E)** Example pressure graph displaying pressure control of iPRS during static pressure myography control. **F)** Difference comparison of P1 and P2 pressure sensors at various set points. The difference begins to drift away as pressure increases beyond the upper limit of the inline sensors of 500 mmHg. **G)** Relationship between duty cycle and flow rate of the bath pump: in our experiments, the duty cycle was set to 80, resulting in a flow rate of 19 mL/min.

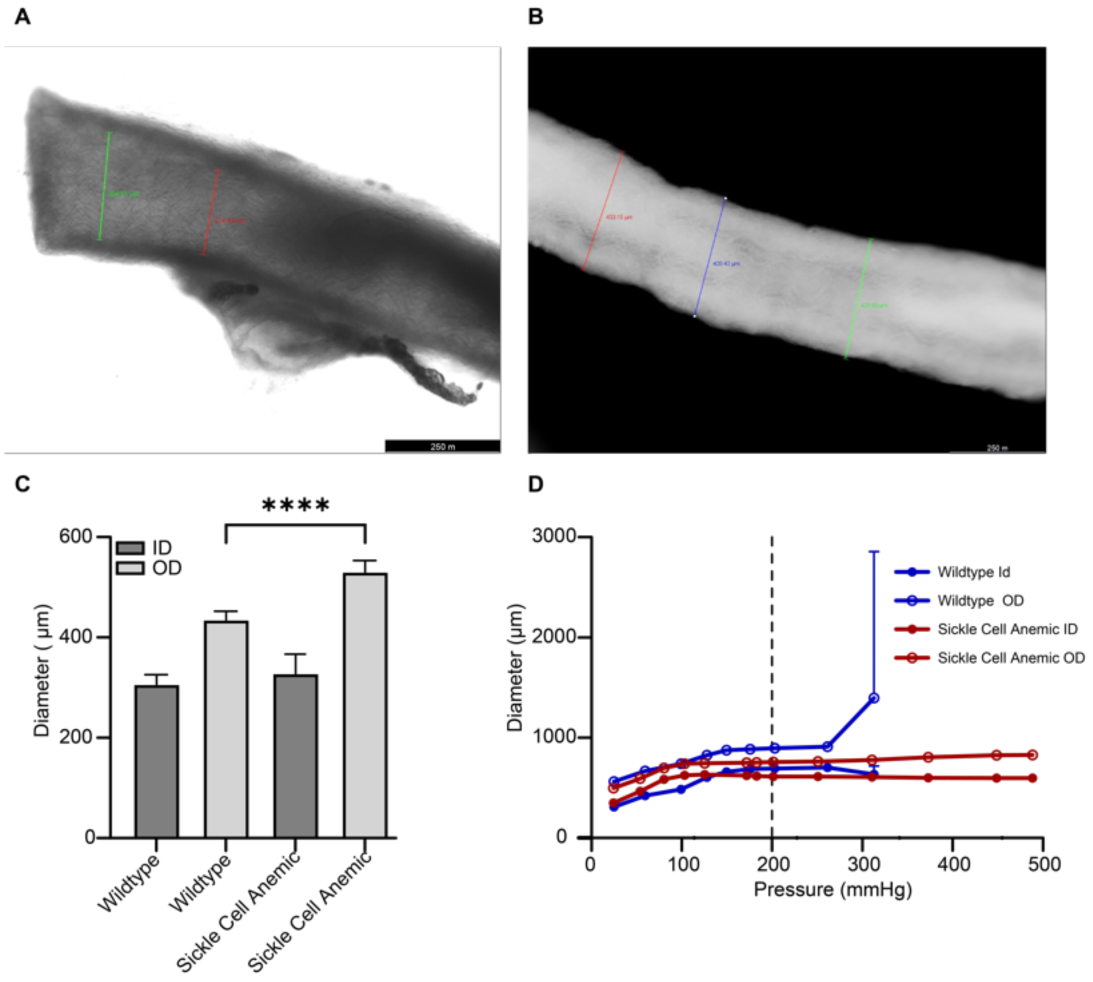


**S2 | A-B)** Bright field and dark field images of Wildtype and Sickle Cell Anemic mouse vessels, respectively. Lecia Measuring tool is super imposed onto the vessels. SB = 250 µm **C)** Comparison of measurements between vessel types, n=5 per measurement, significance measured via student's t-test, *** = p<0.01. Data are represented as mean ± SD **D)** Full static pressure myography graph showing diameter changes of each vessel until maximum pressure measurement allowable of in-line sensors, 500 mmHg.
